# Microfluidic Platform for Automatic Quantification of Malaria Parasite Invasion Under Physiological Flow Conditions

**DOI:** 10.1101/2025.07.29.667357

**Authors:** Emma Kals, Morten Kals, Viola Introini, Boyko Vodenicharski, Jurij Kotar, Julian C. Rayner, Pietro Cicuta

## Abstract

Understanding the impact of forces generated by blood flow on biological processes in the circulatory system, such as the invasion of human red blood cells by malaria parasites, is currently limited by the lack of experimental systems that integrate them. Recent systematic quantification of the growth of *Plasmodium falciparum*, the species that causes the majority of malaria mortality, under a range of shaking conditions has shown that parasite invasion of erythrocytes is affected by the shear stress to which the interacting *P. falciparum* merozoites and their target red blood cells are exposed. Blood flow could similarly impact shear stress and therefore invasion *in vivo*, but there is currently no method to test the impact of flow-induced forces on parasite invasion. We have developed a microfluidic device with four channels, each with dimensions similar to those of a post-capillary venule, but with different flow velocities. Highly synchronised *P. falciparum* parasites are injected into the device, and parasite egress and invasion rates are quantified using newly developed custom video analysis, which fully automates cell type identification and trajectory tracking. The device was tested with both wild-type *P. falciparum* lines and lines in which genes encoding proteins involved in parasite invasion had been deleted. Deletion of Erythrocyte Binding Antigen 175 (PfEBA175) has a significant impact on invasion under flow, but not in static culture. These findings establish for the first time that flow conditions can critically affect parasite invasion in a genotype-dependent manner. The method can be applied to other biological processes affected by fluid motion, such as cell adhesion, migration, and mechanotransduction.

## 1 Introduction

Malaria is a devastating disease responsible for over 600,000 deaths annually, with the majority of deaths caused by *Plasmodium falciparum* infections [1]. The blood stage of the parasite’s life cycle, which causes all the clinical symptoms of the disease, involves parasites invading erythrocytes to form ring stage parasites. The rings develop into schizonts over the subsequent 48 hours, as the parasites develop and multiply inside the erythrocytes. The schizonts then egress, releasing individual parasites known as merozoites that go on to invade new erythrocytes and form new rings [2, 3]. The whole process of egress and invasion typically occurs in less than a minute [4].

There is no direct evidence of where *P. falciparum* invasion occurs in the vasculature. However, postmortem analysis has shown a high level of attachment of infected erythrocytes to capillaries and post-capillary venules in the organs of patients with severe malaria [5]. This points to egress predominantly occurring in the microvasculature, and the subsequent invasion occurring both there and in the post-capillary vessels. However, a merozoite could travel from the narrowest capillary to a large vein within the one-minute invasion window. More recent investigations have also provided evidence that other key sites of invasion are the bone marrow [6] and the spleen [7, 8, 9].

Fluid flow is present in all these environments. Post-capillary vessels have dimensions of roughly 10-40 µm and flow velocities of 0.2-2 mm s^−1^, and the venules that follow have dimensions of 0.1-20 mm and flow velocities of 2-8 mm s^−1^, based on human measurements in the conjunctival pre-capillary arterioles of the eye [10, 11] and in human fingers [12]. This fluid flow generates shear forces that could impact the interaction between merozoites and red blood cells and, therefore, affect invasion.

Despite invasion occurring under dynamic flow conditions *in vivo*, most parasite growth and invasion assays are conducted in static *in vitro* conditions, with blood sedimented to the bottom of tissue culture flasks or plates. It has been shown that fluid flow generated by culture on an orbital shaker can both enhance growth rates [13, 14, 15, 16, 17] or reduce growth rates [18]. In our recent study, we found that at shear forces comparable to those measured previously in the microvasculature, growth was reduced across wild-type lines, with the magnitude of the effect greater in lines missing key proteins known to be involved in the early steps of erythrocyte attachment [19]. Growth plates on orbital shakers are, however, not good mimics of *in vitro* flow conditions, as they generate bulk swirling rather than vessel-like wall shear stress [20].

Many microfluidic devices have been developed to mimic the conditions of the circulatory system, as reviewed by [21]. Microfluidic devices have also been built to explore other aspects of malaria biology, such as infected erythrocyte deformability [22, 23, 24], sorting of infected erythrocytes from non-infected [25, 26] and investigating the cytoadhesion of infected erythrocytes to endothelial proteins or cells [27, 28, 29]. However, no microfluidic device has been developed to follow malaria invasion under flow conditions.

We have developed a microfluidic setup that allows the invasion of *P. falciparum* into human erythrocytes to be tracked under controlled flow conditions. We tightly synchronise our *P. falciparum* parasites before loading the device so that most schizonts egress within a few minutes, resulting in egress and invasion occurring as the sample flows through parallel microfluidic channels under controlled flow. We also developed an image analysis pipeline that enables the automated identification and tracking of different cell types, allowing us to quantify the invasion rate.

Using this platform, we explored how physiologically relevant flow influences parasite invasion. Our experiments revealed that while the invasion phenotype of wild-type parasites is not impacted by the range of flow rates investigated, disruption of one invasion ligand, PfEBA175, but not another, PfRH4, can reduce invasion under higher flow rates. The uncovering of a novel flow-based phenotype highlights the value of this microfluidics-based assay for investigating parasite–host interactions under controlled flow conditions.

## 2 Results

### 2.1 Design of parallel-flow microfluidic device for erythrocyte invasion assay

The PDMS microfluidic device we developed comprises four parallel channels, which can each be set to have a different flow rate; the design is illustrated in fig. 1. The chip has two inlets, one for loading and one for driving the flow from a piezoelectric disc pump. The pump maintains a constant inlet pressure using a feedback controller, set to a 190-240 mbar. The inlet then branches into four parallel channels with identical dimensions. Each channel is 6.7 µm high and 27.7 cm long. To allow us to observe egress and invasion in the channel, at least a 10 min time window is required, which limits the maximum flow velocity to 460 µm s^−1^. The channel depth of 6.7 µm ensures that there is a single layer of erythrocytes, which is necessary to allow automated detection of erythrocytes in brightfield. Channels deeper than 6.7 µm were tested but resulted in out-of-focus erythrocytes and multiple layers of erythrocytes, which hindered automated cell tracking, fig. S1. Each channel has an outlet connected to a vertical water column of varying height, which generates different backpressures and results in a distinct flow velocity, allowing four different velocities to be compared simultaneously in the same chip run. There are valves on the water column which are set to block the water column when loading, ensuring rapid loading for all channels. The valves are then switched after loading to connect the water columns. The channels are 100 µm wide, meaning that all four channels fit into a single field of view with the 20x objective. Observations are made at a single point near the outlet (imaging region), and the same imaging region is continuously monitored throughout the experiment, fig. 1a.

**Figure 1.**
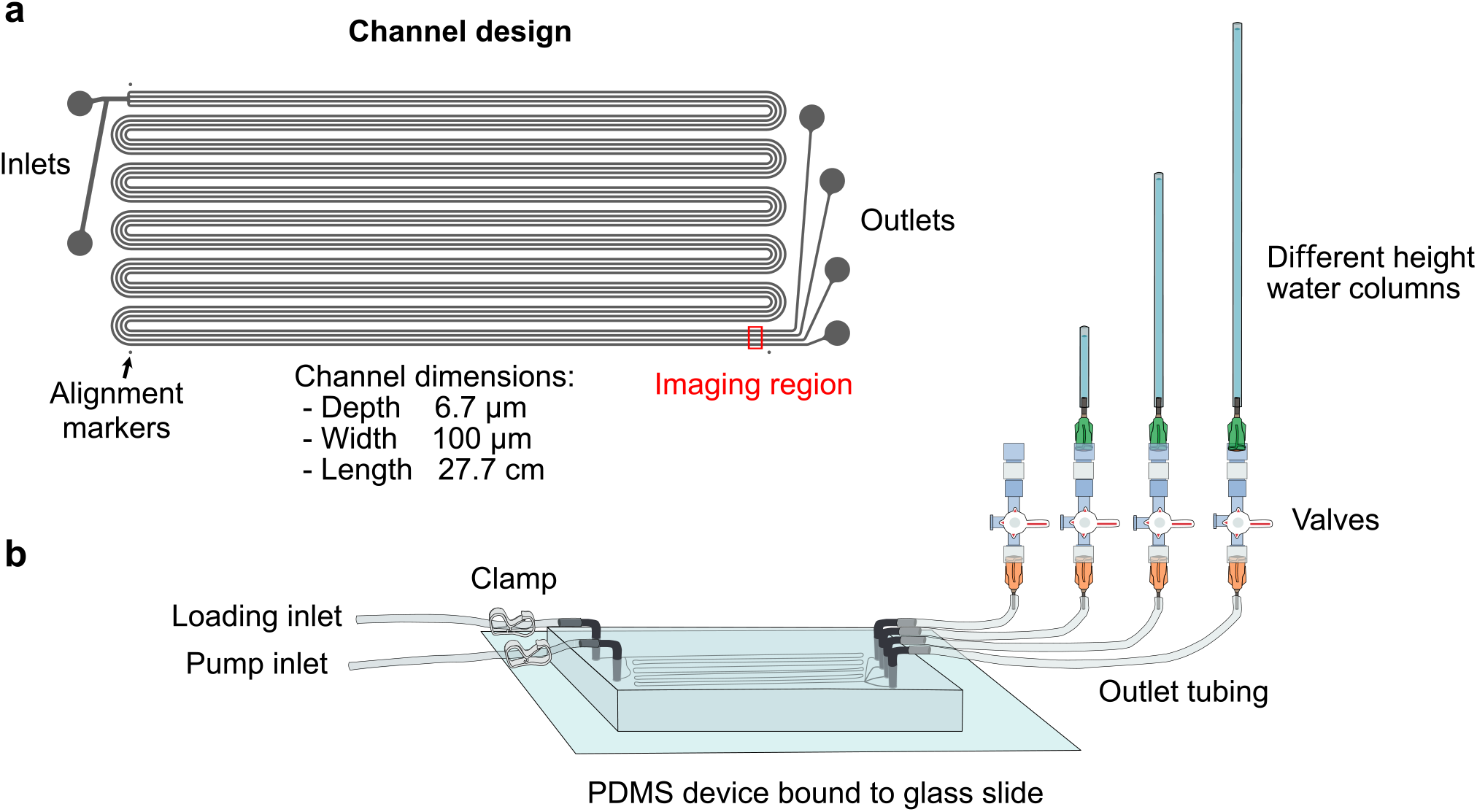
A PDMS chip has been designed to enable parallel measurements of cells flowing in four different wall shear profiles. **(a)** Shows the 2D channel design, which is printed onto photoresist on a silicon wafer with a height of 6.7 µm. This is then cast in PDMS to create the chip. **(b)** Shows how the tubing is attached to the PDMS devices. One inlet allows loading, and the other is connected to a pressure pump that drives the flow. There are clamps on the inlet tubing to control which is connected to the device. The inlet channel branches into four channels that snake back and forth, each with its own outlet. Each outlet is connected to a water column of different heights that generates backpressure, with the result that each channel has a different flow velocity. There are valves on the water columns that can block the channels and allow rapid homogeneous loading of all the channels.

In this platform, the experimentally controlled variable is the velocity of cells in each channel, quantified directly by particle tracking in the imaging region, fig. S8. Quantities such as wall shear stress and hydrodynamic force are not measured directly but are inferred from the measured velocity using a laminar-flow model for the channel geometry, as outlined in Section S1. Because shear forces vary spatially across the cross-section of the channel, we report inferred values as maximum shear stress rather than as precise shear stresses experienced by every cell.

### 2.2 Tight synchronisation of *P. falciparum* parasites ensures a high invasion rate in flow

To facilitate a high invasion rate within a short window of time, the parasites were highly synchronised through multiple developmental cycles (see Methods), then treated with Compound 2 for 3-5.5 hours. Compound 2 is a *P. falciparum* protein kinase G inhibitor which blocks schizont egress, resulting in the accumulation of highly mature schizonts pre-egress [30]. Once Compound 2 is removed, the parasites will egress approximately 15 minutes later. As washing, transfer to the microscope, and loading take around 10 minutes, the schizonts typically start egressing within 5 minutes of the channels being loaded. This protocol ensures that the majority of parasites are at the same very late schizont stage when loaded into the device and will start egressing quickly once loaded. This maximises the number of invasion events observed during each experimental run, and allows a well-defined timeline in the analysis.

### 2.3 Automated cell identification and tracking

The channels are imaged at a single imaging field of view close to the outlet, referred to as the imaging region, fig. 1a. The sample is initially homogeneous throughout the device; therefore, the cell counts in the first few minutes, before egress, will be the same in all channels. The parasites then begin to egress and invade, as they flow through the device, fig. 2a. The flow rate determines how long it takes for cells that would have been in the inlet at the start of the experiment to reach the imaging region. That time interval is the longest that we include in our analysis, thus ensuring that all the invasions measured will have occurred in the channel under the measured steady flow rate.

**Figure 2.**
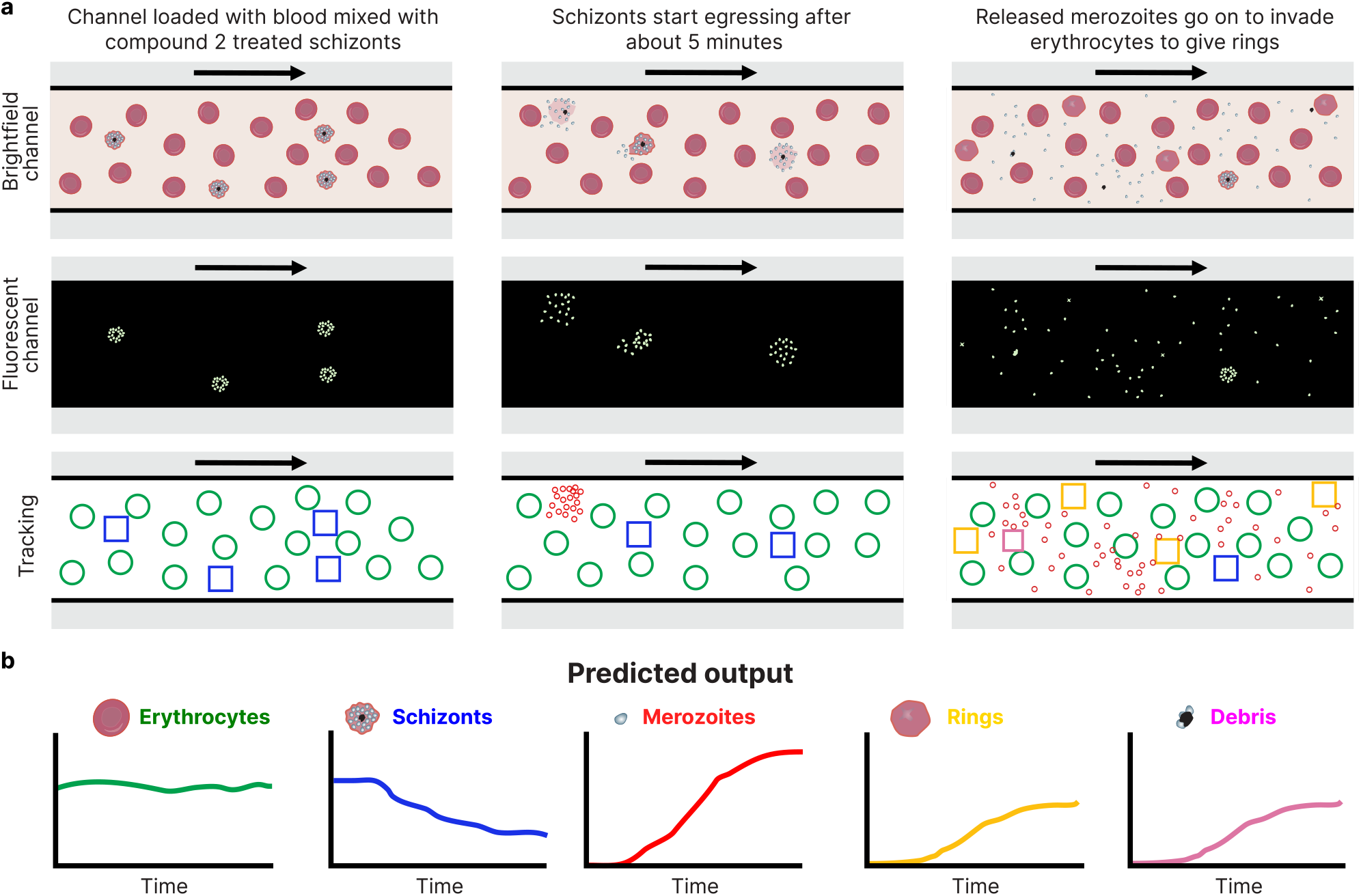
Experiment design. **(a)** Cartoon to show how the different cell types are identified throughout the experiment in our imaging region, which is close to the outlet. The experiment starts when the sample is injected into the channel. To begin with, throughout the channel, only erythrocytes (green circles) and schizonts (blue squares) are present. Then, after *∼* 5 minutes, the schizonts start to egress throughout the channel, releasing the merozoites (red circles). By the end of the video, some of the released merozoites have invaded erythrocytes and formed rings (yellow squares). **(b)** Trends of the predicted changes in the four different cell types, and cell debris. Invasion rates are determined by looking at the change in cell count between time window 1 (W1), highlighted in pink shading, and time window 2 (W2) in blue shading. A vertical line represents the time point at which cells that started at the inlet port at time zero pass through the imaging field of view, the position of which is determined by integrating the flow velocity of the cells in each channel. No data is used after this time point, as any invasion that occurs after this line could have occurred before the cells entered the microfluidic channel.

In order to quantify invasion, we have automated the process of cell type identification and velocity tracking. Videos of the channels are recorded in both brightfield and fluorescence. Automated detection was set up to identify erythrocytes, schizonts, merozoites, rings, and cell debris. Cells are tracked as they pass through the 224 µm of the imaging region. Erythrocytes were detected using brightfield images based on cell size and shape, while parasite stages were stained with the DNA dye SYBR Green, which labels parasite nuclei but not erythrocytes, as they are anucleate. Schizonts were distinguished from rings and merozoites by their large, bright fluorescent signal, reflecting the multiple parasites and therefore multiple copies of genomic DNA (gDNA) they contain. In contrast, rings and merozoites each contain a single copy of gDNA and therefore exhibit weaker, dimmer fluorescence. Examples of the different cell types are shown in fig. 3. To distinguish rings (parasites that have invaded a new erythrocyte) from free merozoites that happen to visually overlap with an erythrocyte as they are both carried along by flow, we scored rings only where an erythrocyte had a weak fluorescent signal that travelled with it across the entire imaging window. Sometimes, when schizonts rupture, in addition to free merozoites, a clump of merozoites or other membrane debris is released. We therefore quantified an additional category, called ‘cell debris’ (fluorescent material of intermediate intensity), to ensure that we did not count these clumped parasites, which are unlikely to successfully invade, as either schizonts or merozoites.

**Figure 3.**
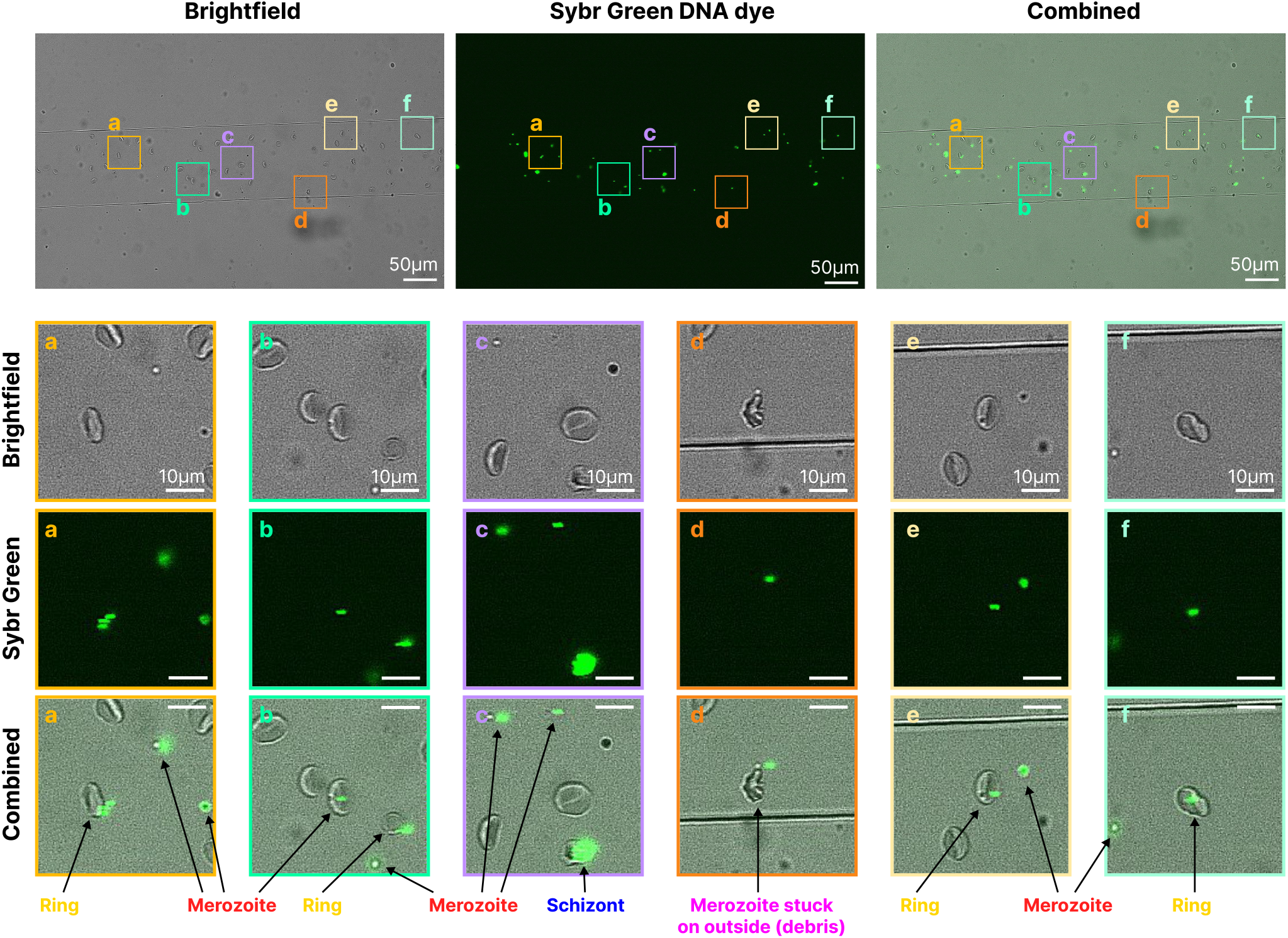
Examples of brightfield and fluorescent images with annotated cell classification. Infected erythrocytes stained with SYBR Green at a concentration of 1 in 5000 for 45 minutes. Brightfield and fluorescent images are captured sequentially, making the moving cells appear slightly offset. Callouts **a, b, e**, and **f** show one ring each, meaning that when the erythrocytes are followed across the channel, the fluorescent signal stays with them. **b** shows a merozoite above an erythrocyte, but the fluorescent signal does not follow the erythrocyte over time, indicating this is not a ring and underlining the importance of tracking the species over time for correct classification. **c** shows an example of a schizont, with a much stronger fluorescent signal. **d** shows an example of a merozoite stuck on the outside of an erythrocyte, which is classified as debris.

With this experimental design, we expect to see: (1) an almost constant population of erythrocytes, since the number of infected erythrocytes in any parasite culture is low relative to the overall number of erythrocytes (usually <5 %). However, given the tight synchrony of the starting parasites, we should expect a (2) drop in the population of schizonts 5-10 minutes after injection into the devices, as they mature and rupture after Compound 2 has been removed, and then (3) a subsequent increase in the number of free merozoites (from a near zero starting population) as they are released from the egressing schizonts. Finally, (4) as the released merozoites invade erythrocytes, the population of rings should increase slightly later than when merozoites first appear, fig. 2b.

### 2.4 Automated tracking was able to identify different cell types with a high accuracy rate

We carried out at least four biological replicates (each performed with parasite samples prepared on different days and with blood from different donors) for each of two wild-type *P. falciparum* lines (NF54 and 3D7). We first visually confirmed that egress had occurred and that released merozoites invade erythrocytes to produce rings, fig. S6. We then assessed tracking accuracy by manually annotating cell types in selected frames and comparing them to the automated analysis, which demonstrated a high level of accuracy. Table 1 shows the overall accuracy based on manually labelled images from four experiments and three time points (1, 5 and 8 minutes after invasion). For each sample frame, the number of erythrocytes, merozoites, rings, schizonts and cell debris present in each channel was manually counted. This was then compared to the mean cell counts per channel reported by the automated analysis at the timestamp of that frame. The error rate is calculated as the difference between the manual and automated cell counts divided by the manual cell count. If both counts are zero, the error is set to zero. If the manual count for a cell type was zero, the error for that repeat was excluded from the average. See Table S1 for the full list of error rates.

**Table 1:** Average error rates comparing manual and automated counts at a given time point. Time points after 1, 5 and 8 minutes of invasion. The error rates were calculated by comparing frames in which the different cell types were manually counted to the counts for the same frames that were determined using the automated analysis code.

| Parameter | Erythrocytes | Schizonts | Merozoites | Rings | Cell debris |
| --- | --- | --- | --- | --- | --- |
| Error rate | 16% | 34% | 4% | 14% | 51% |
| Total counted | 2322 | 57 | 247 | 150 | 15 |

Next, we explored how the population of each cell type changed over time for each channel. An example is shown in fig. 4, with similar plots for the remaining experiments in fig. S7. Broadly, the populations changed as expected (fig. 2b), with an initial reduction in schizonts followed by an increase in merozoites and rings, confirming that the automated cell tracking method was detecting the sequential biological steps of schizont rupture, merozoite egress and invasion. Tracking the average speed of all erythrocytes across channels confirmed that the backpressure approach resulted in different flow velocities, fig. S8. Repeat 2 of NF54 for example has flow velocities ranging from 35 µm s^−1^ in channel 1 to 320 µm s^−1^ in channel 4. fig. S8 confirms that the flow velocities of the different cell types are also highly consistent, indicating that the size or other properties of the cells do not generally affect their speed. There are some notable exceptions, where cells, most often merozoites, become adhered to the glass coverslip for either very short or longer durations, resulting in a temporary drop in the recorded speed for that cell category.

**Figure 4.**
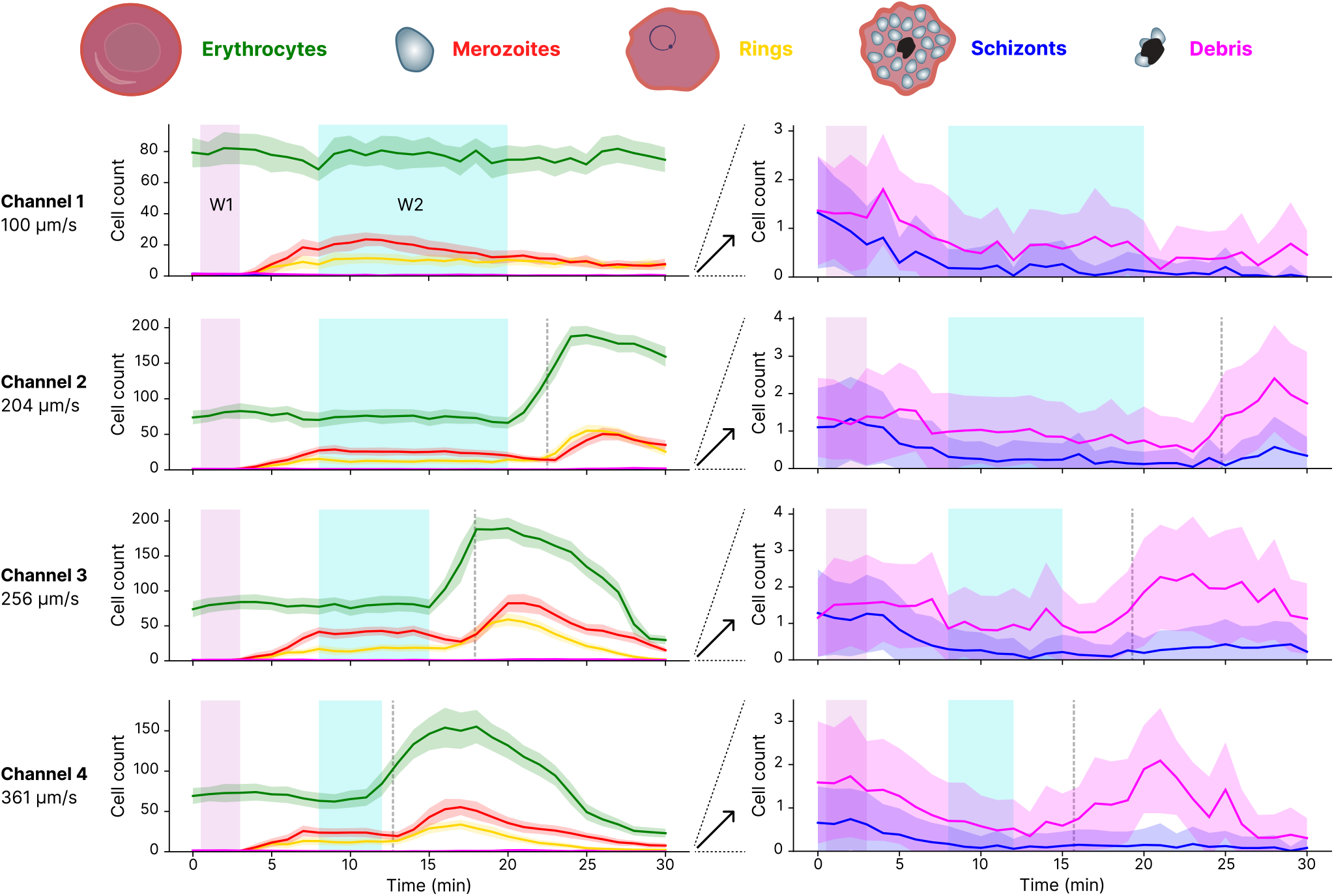
Example from a single experiment of tracking the populations of cells in each of the four channels. The data shown is for a single experiment run with the wild-type line NF54. Cell count refers to the number of cells we count in our field of view for a given channel at a given point in time, averaged over a 30 s interval. This averages all the cells that are tracked as they flow through the imaging window in the 30 s window. The graphs on the right show the same data as on the left but for a smaller cell count range, to visualise what happens with the schizont and cell debris populations where there are many fewer individual cells. The different colours show the different cell types: green is erythrocytes, red is merozoites, yellow is rings, blue is schizonts, and pink is debris. The time windows are highlighted with window 1 (W1) highlighted in pink shading and window 2 (W2) in blue shading. The windows are used to quantify the change in cell populations. A vertical dashed line is added at the time point where cells that were at the inlet port at time zero pass through the imaging field of view, the position of which is determined by the flow velocity of cells in the channel. No data after the dashed line is analysed. Blood settles in the inlet after loading, causing a wave of higher erythrocyte density to reach the imaging window. However, as this blood is in the inlet at the start of the experiment, it is always after the cut-off point (dashed line) at which we stop using the data for a channel, and so it does not affect the results. The velocities shown for each channel represent the mean velocity of all the cells tracked within that channel.

### 2.5 Invasion efficiency was quantified by calculating the proportion of rings

The invasion rate can be calculated by measuring the relative change in the proportion of rings in the imaging region between two time windows: an early window 1 (W1), which provides the cell count just after loading and before schizonts begin to egress, and a later window 2 (W2), which provides the cell count after the bulk of successful invasion events have occurred. The default settings for W1 were between 0.5 and 3 minutes after the start of the experiment, and W2 was between 8 and 15 minutes. Window 1 is started after at least 30 seconds to allow the system to stabilise, and the flow velocities to settle after the experiment is started. The end of W1 and the beginning of W2 were chosen based on observations of the video files and cell count traces over time, and the selection of points where the transition from pre-egress to post-invasion occurred. The timing of egress is fairly consistent, as it is determined by the time since Compound 2 was removed. The end of W2 was set to 15 min as default, but was adjusted on a channel by channel basis to a) consider the flow velocity to ensure only parasites that were in the device at the beginning of the experiment are included, b) not include any significant changes in haematocrit and c) last for longer if the flow velocity is very low and the cell counts are stable to improve statistical accuracy. By adjusting the time window based on the flow velocity and channel length, we ensure that only egress and invasion that occur under flow in the channel are considered, and not events that occurred in the inlet before they are exposed to flow conditions.

The change in cell count for each cell type can then be calculated by subtracting the mean count in window 2 C_*W* 2_ from the mean count in window 1 C_*W* 1_, namely ∆C = C_*W* 2_ − C_*W* 1_. For example, we compute the proportion of rings as:

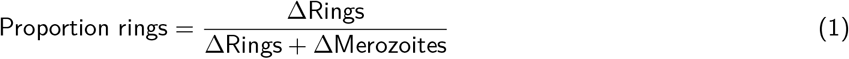

*Proportion of rings* was the main metric we used to measure the invasion rate; it effectively captures the number of egressed merozoites that have successfully invaded erythrocytes. For all experiments we also calculated invasion per schizont as Invasions per schizont = − ∆Rings*/*∆Schizonts, but we found this metric had a low signal to noise ratio due to the small number of schizonts present in each experiment (often less than one visible in the field of view at any point in time, as opposed to tens of rings and merozoites in window 2).

### 2.6 Invasion efficiency for 3D7 and NF54 shows no significant difference under flow

We assessed the invasion of wild-type lines NF54 and 3D7 under different flow velocities. The NF54 strain is most likely of West African origin [31], and 3D7 is a clone of NF54 [31] that has been cultured independently for 30 years and, while being genetically very similar [32], has multiple diverged phenotypes [33, 34, 35]. As each run resulted in slightly different flow velocities, we grouped the data into low (27-150 µm s^−1^), medium (150-300 µm s^−1^) and high (300-410 µm s^−1^) flow rates. Scatter plots are also shown in fig. S9. Across the flow velocities tested, there was no significant difference between the lines for any of the flow ranges. There was also no significant difference in the proportion rings at any flow range between the two strains, fig. 5a.

**Figure 5.**
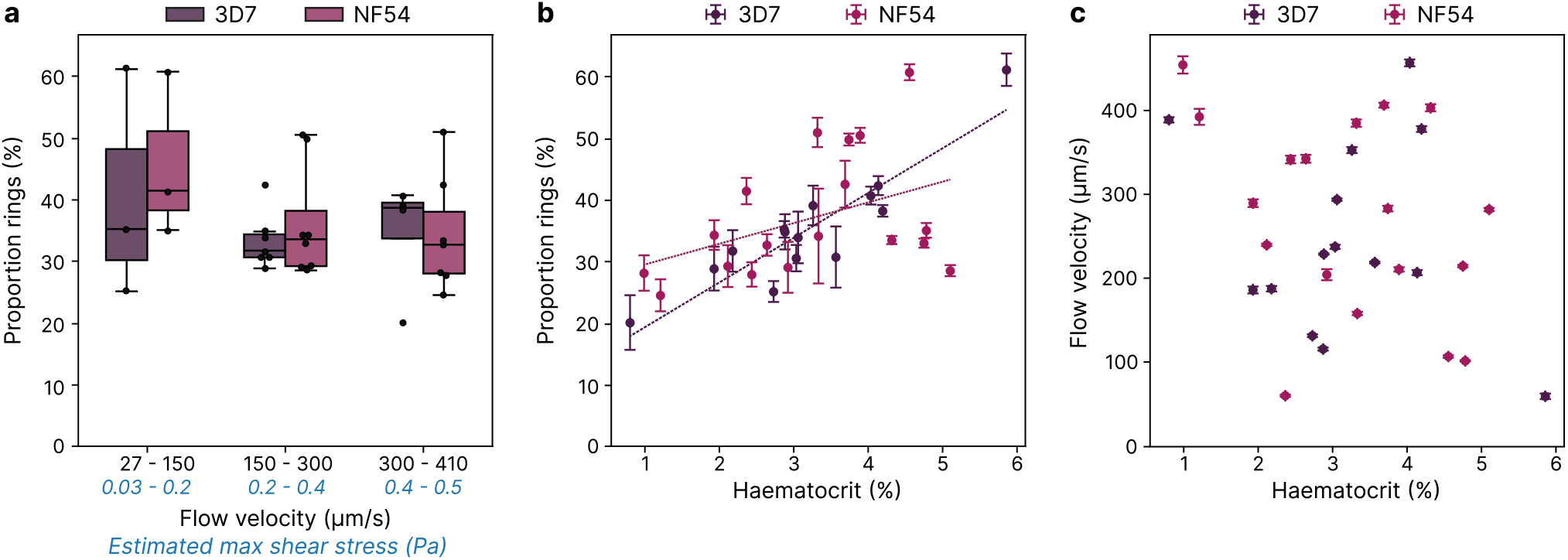
There is no significant difference in invasion with flow velocity but a positive correlation between invasion rate and haematocrit for wild-type lines NF54 and 3D7. The data shown is for wild-type lines of NF54 and 3D7. **(a)** Boxplot showing how invasion rate, measured as the proportion of erythrocytes that have invaded to form rings (*proportion rings*), is affected by flow velocity. Flow velocity was binned to low (27-150 µm s^−1^), medium (150-300 µm s^−1^) and high (300-410 µm s^−1^). Each data point represents one channel in one experiment run, with three to eight points per condition. None of the groups showed statistically significant differences as determined by t-tests performed at the 5% significance level (p *<* 0.05). The estimated maximum shear stress present in the microfluidic channels at each flow velocity (calculated in Section S1) is provided for reference. **(b)** A scatterplot of proportion rings vs blood haematocrit is shown. Haematocrit is measured based on the number of erythrocytes observed and the volume of the observed channel. Each data point represents data from one channel and one experiment, with error bars in x and y representing standard deviation. Least squares linear regression is shown with slope *m* and coefficient of determination *R*2 shown for each strain. **(c)** The scatterplot of flow velocity vs haematocrit shows how these are independent variables in our experimental setup with no apparent correlation.

### 2.7 Invasion rate is dependent on haematocrit

Next, we assessed whether invasion was impacted by other factors such as haematocrit, initial parasitaemia, average flow velocity, invasion per schizont, proportion rings and the egress rate, see fig. S9. We found the most noteworthy relationship to be a positive correlation between haematocrit (erythrocyte density) and invasion rate, fig. 5b. Experiments were run with the sample loaded at 10% or 15% haematocrit, but the resultant haematocrit was measured to be between 1-6%, and even between channels of the same device, the measured haematocrit varied. We are not sure of the cause of this variation, but it may be due to variability in the efficiency of blood entering the narrow channels. The haematocrit within a given channel was consistent from the start of time window 1 to the end of time window 2, varying by 11 *±* 6 % (min of 3.9% and max of 32%) across all experiments. Haematocrit in the imaging region increases just before cells that were at the inlet port at time zero pass through the imaging field of view, as erythrocytes that had sedimented in the inlet reach the imaging field. This, however, was never included in the analysis as time window 2 was always set before this increase. For the wild-type lines, the correlation between haematocrit and proportion rings was weaker for NF54 (slope=3.3 and R2=0.17) and stronger for 3D7 (slope=7.2 and R2=0.82). This correlation is not unexpected because the higher the haematocrit, the greater the probability of a merozoite colliding with an erythrocyte while it is still invasion-competent. No correlation was seen between haematocrit and flow velocity, fig. 5c, suggesting that density of erythrocytes does not impact flow. This scatter also indicates that the relationships presented in fig. 5a are not just a side effect of systematic variations in haematocrit.

### 2.8 A *P. falciparum* strain lacking the PfEBA175 invasion ligand invades less efficiently under higher flow rates

We had previously observed that conditions of high wall shear stress, created by orbital shaking, cause a greater reduction in growth in several genetically modified *P. falciparum* lines in which members of the PfEBA and PfRH invasion ligand family had been individually deleted [19]. Critically, as reported by both us and others, under static conditions these lines grow at the same rate as the wild-type line they are derived from [36, 37, 38, 39, 40, 41, 42]; the invasion defect is only revealed when shear stress is added [19], presumably increasing the forces that oppose merozoites and erythrocytes making productive cell-cell contacts. To explore whether flow conditions could have the same effect, we tested two of these lines using our microfluidic flow approach: ∆PfRH4, in which we had previously seen a weaker effect of shaking on invasion, and ∆PfEBA175, which had the largest reduction in growth under shaking [19]. The invasion efficiency of ∆PfRH4 was not significantly different between low, medium and high flow velocities, just like the wild-type strain from which it was generated, NF54. By contrast, while ∆PfEBA175 invasion was not significantly different to the other lines at a low flow velocity, it was significantly reduced at both medium and high flow velocities, with the proportion of ring-stage parasites reduced more than 2-fold relative to wild-type at the highest flow rate. ∆PfEBA175 also showed a significantly lower invasion rate relative to itself at low flow velocities compared to medium and high flow velocities, fig. 6. This indicates that the invasion of lines lacking PfEBA175 is more impacted by flow conditions in the 150-410 µm s^−1^ range than PfRH4, suggesting that PfEBA175 plays a particularly critical role in merozoite-erythrocyte adhesion under physiological flow conditions. As for the wild-type lines, there is a positive correlation between haematocrit and invasion rate, fig. S10a. Again, there was no correlation between haematocrit and flow velocity, fig. S10b.

**Figure 6.**
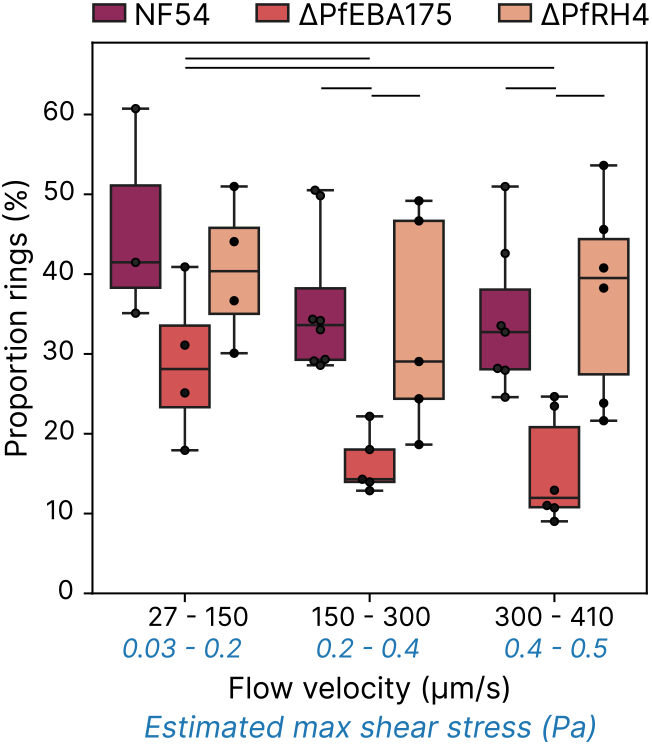
Increasing flow velocity significantly reduces the invasion rate of ∆PfEBA175. ∆PfEBA175 and ∆PfRH4 lines are constructed in the NF54 background. Boxplot showing how invasion rate, measured as the proportion of erythrocytes that have invaded to form rings (*proportion rings*), is affected by flow velocity. Flow velocity was binned to low (27-150 µm s^−1^), medium (150-300 µm s^−1^) and high (300-410 µm s^−1^) with estimated max shear forces generated by this flow included. The significance lines indicate statistically significant differences between groups, as determined by t-tests performed at the 5% significance level (p *<* 0.05). Each data point represents one channel in one experiment run, with three to eight points per condition. The estimated maximum shear stress present in the microfluidic channels at each flow velocity is calculated in Section S1.

### 2.9 Flow rate does not impact contact probability between merozoites and erythrocytes

Finally, we explored whether the changes in invasion rates observed under different flow conditions for ∆PfEBA175 could be explained by changes in the contact probability between merozoites and erythrocytes. Across all experiments, there was generally little difference between the mean velocities of red blood cells (RBCs; here defined as erythrocytes, schizonts, and rings) and merozoites in a given channel, fig. 7a. We next quantified, for each channel, the proportion of merozoites that contacted an RBC within the imaging region. Merozoites that contacted RBCs tended to have a slightly lower mean velocity than the overall merozoite population, fig. 7b. This is consistent with the idea that merozoites moving more slowly, for example, near channel walls, have more opportunity to encounter RBCs. However, when we compared the proportion of merozoites that contacted RBCs to the mean merozoite velocity across the full range of flow conditions tested (27–410 µm s^−1^), we did not observe any correlation, fig. 7c. These findings show that flow rate does not modulate the likelihood of merozoite–erythrocyte encounters, implying that the flow-dependent reduction in ∆PfEBA175 invasion arises from the hydrodynamic forces experienced during attachment rather than from any reduction in contact opportunities.

**Figure 7.**
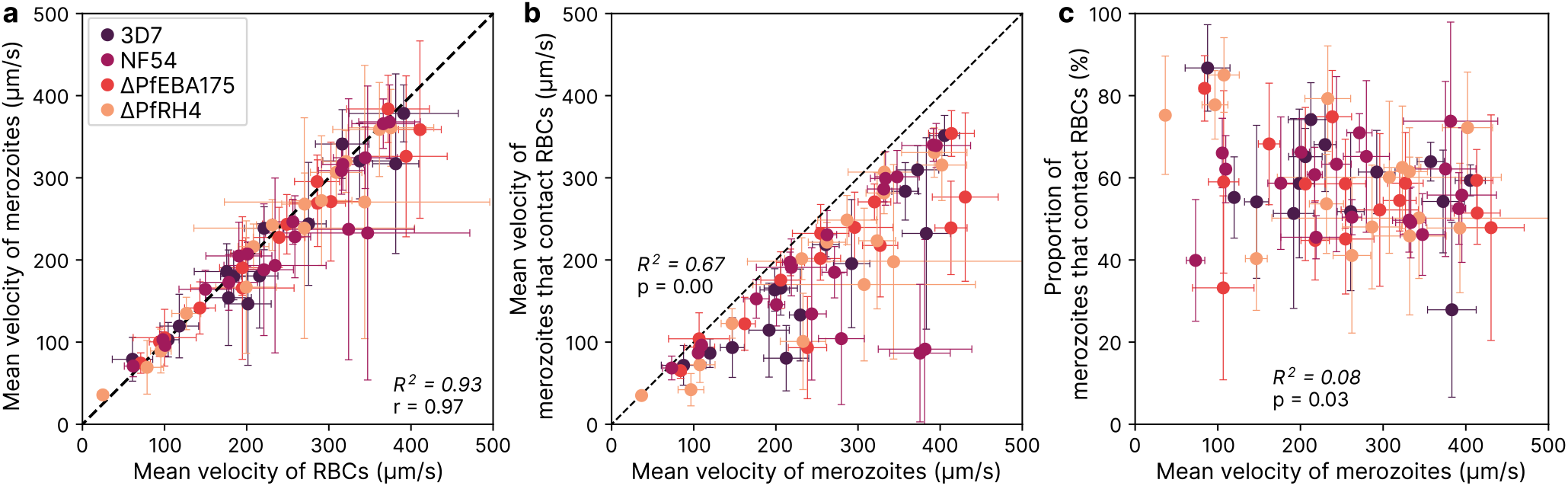
Flow velocity does not meaningfully impact contact probability in this flow range. **(a)** Scatterplot showing how merozoites and RBCs move at roughly the same speed through the channels. Each point corresponds to the data from one channel between the start of window 1 and the end of window 2, with error bars representing standard deviation. **(b)** Shows that the merozoites that collide with RBCs generally move a bit slower than the ones that don’t. This suggests that if a merozoite randomly moves slowly (because it is very close to a wall for example) it is more likely to collide with RBCs moving past. **(c)** The proportion of merozoites contacting RBCs (calculated as 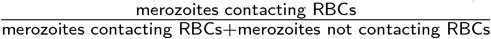) does not depend on flow rate.

### 2.10 The hydrodynamic forces experienced by merozoites during invasion are around 10 pN

We estimated the forces acting on merozoites in Section S1, where we show that the flow in our device operates at Reynolds number *≪* 1. This places the dynamics firmly in the creeping-flow (Stokes flow) regime, in which inertial forces are negligible and viscous forces dominate. We then estimated the hydrodynamic load on a merozoite by calculating both the shear stress acting on its surface and the Stokes drag experienced in a linear shear field. These independent approaches yield consistent force estimates of approximately 10 pN. This value represents the characteristic load on a merozoite that is attached to an erythrocyte near the channel wall under the flow conditions used. The forces experienced by freely moving merozoites will generally be lower, and transient interactions (for example collisions with passing RBCs) may generate larger instantaneous forces.

## 3 Discussion

Our study demonstrates a novel microfluidic assay that enables direct quantification of *P. falciparum* invasion under defined laminar flow conditions that overlap with the lower end of reported microvascular conditions. The invasion phenotype of the NF54 and 3D7 wild-type lines and ∆PfRH4 are not affected by these flow conditions, but ∆PfEBA175 has a clear reduction in invasion efficiency as flow rates increase, fig. 5a and fig. 6. This shows how flow and shear can reveal invasion defects that are missed by static assays. Further experiments, including greater flow ranges, more replicates and less variation in haematocrit, would be required to precisely elucidate the relationship between flow and invasion without the need for binning.

Our calculations show that, in the 27-410 µm s^−1^ range tested here, the reduction in ∆PfEBA175 invasion efficiency is likely driven by hydrodynamic forces opposing attachment rather than changes in contact probability between erythrocytes and merozoites. We estimate that merozoites are experiencing up to 10 pN of hydrodynamic load at our maximum flow velocity of 410 µm s^−1^. Previous measurements using optical tweezers showed that detaching ∆PfEBA175 merozoites from erythrocytes requires forces in the range 12-47 pN [36]. Because these measurements were performed after giving the merozoites several seconds to attach, the force needed to prevent attachment is expected to be lower. Therefore, a hydrodynamic load on the order of 10 pN opposing attachment provides a consistent explanation for the reduced invasion efficiency of ∆PfEBA175 at higher flow rates. Notably, the highest flow rate tested here produces a shear stress of 0.5 Pa, which is lower than the wall shear stress (WSS) measured in the human conjunctival capillaries of 1.5 Pa [11]. Therefore, the hydrodynamic force opposing invasion *in vivo* could be even stronger than that tested here, which we speculate could mean that PfEBA175 may be even more important in maintaining attachment under flow *in vivo*. It is also important to note of course that the *in vivo* environment will be much more complex than this model, where the presence of endothelial cells will make flow rates and shear stress much more heterogeneous, and where flow will be pulsatile rather than constant. While we believe that this design is a step towards understanding invasion *in vivo*, there is a clear need for caution in interpretation.

This raises the question of why high flow rates only reduced invasion efficiency in the ∆PfEBA175 line. This outcome is especially striking, considering that our previous optical-tweezer measurements showed a significantly lower detachment force for ∆PfRH4, whereas ∆PfEBA175 did not differ from the NF54 wild-type background. PfEBA175 binds to glycophorin A (GYPA) [43, 44], which is the most abundant glycophorin protein on the surface of erythrocytes with one million copies per cell [45]. PfRH4 binds to Complement C3b/C4b receptor 1 (CR1), a type 1 membrane glycoprotein which is expressed at far lower levels of approximately 50-1200 molecules per erythrocyte [46]. This leads us to speculate that PfEBA175 may play a particularly important role in the earliest stages of attachment. In its absence, initial contacts may be weaker and so less able to withstand flow-generated hydrodynamic forces, leading to reduced invasion despite normal detachment forces measured after several seconds of binding. In contrast, PfRH4 may alter binding strength at a later stage of invasion, thereby affecting measured detachment force without substantially altering early attachment under flow.

∆PfEBA175 being the most affected by flow conditions is consistent with our previous measurements under high WSS generated by culturing parasites on an orbital shaking platform. In those experiments, ∆PfEBA175 had the largest reduction in growth rate relative to static conditions among all PfEBA and PfRH knock-out lines tested [19]. Table 2 summarises the invasion phenotypes observed in this shaking assay and compares them with the results presented in this paper. It is important to emphasise that these assays are very different, so direct comparisons must be made with caution.

**Table 2:** Comparing results from this microfluidics assay with our previous growth assay performed on orbital shakers [19]. All values are reported as mean *±* standard deviation, with flow conditions, estimated max wall shear stress (WSS), and estimated max force experienced by a merozoite during invasion reported for comparison. Creeping flow is a type of laminar flow where inertial forces are negligible. Invasion rate is defined as the increase in rings divided by the reduction in schizonts over a 3.5-hour window. See section S1 for force estimates. Microfluidic flow ranges are low (27-150 µm s^−1^), medium (150-300 µm s^−1^) and high (300-410 µm s^−1^), with estimated WSS and forces based on the average velocity in the range. The shaking and microfluidics assays were performed concurrently, using samples from the same synchronisation cycles.

| Strain | Proportion rings (%)<br>from microfluidics assay |  |  | Invasion rate (ratio)<br>from shaking assay |  |  |  |
| --- | --- | --- | --- | --- | --- | --- | --- |
|  | Low flow | Medium flow | High flow | Static | 45 rpm | 90 rpm | 180 rpm |
| 3D7 | $40 \pm 20$ | $33 \pm 4$ | $30 \pm 10$ | $9 \pm 3$ | $12 \pm 2$ | $11 \pm 2$ | $17.7 \pm 0.2$ |
| NF54 | $50 \pm 10$ | $36 \pm 9$ | $34 \pm 9$ | $11 \pm 4$ | $9 \pm 4$ | $6.9 \pm 0.3$ | $15 \pm 2$ |
| $\Delta\text{PfEBA175}$ | $30 \pm 10$ | $16 \pm 4$ | $15 \pm 7$ | $8.2 \pm 0.8$ | $10 \pm 3$ | $4.0 \pm 0.9$ | $13 \pm 1$ |
| $\Delta\text{PfRH4}$ | $40 \pm 9$ | $30 \pm 10$ | $40 \pm 10$ | $10 \pm 4$ | $10 \pm 3$ | $6.5 \pm 0.9$ | $19 \pm 5$ |
| **Flow conditions** | creeping | creeping | creeping | none | laminar | laminar | turbulent |
| **Estimated WSS (Pa)** | 0.1 | 0.4 | 0.5 | 0 | 0.63 | 2.5 | unknown |
| **Estimated force (pN)** | 1.6 | 3.4 | 5.7 | 0 | 7.9 | 31 | unknown |

This microfluidic assay has been developed to assess invasion under physiological flow conditions and is not intended to replace traditional invasion assays, as it is more technically challenging and has lower throughput. Traditional growth and invasion assays require no specialised consumables and can be quantified by flow cytometry, enabling measurements in a 96-well plate format. If a panel of genes were to be tested for their impact on invasion under high-shear stress conditions, we envisage screening them first using a high-throughput growth assay under static and orbital shaking conditions. Then the lines of interest could be tested in the microfluidic assay to validate their impact under more physiological conditions.

Further development of this assay could address many other aspects of how flow impacts invasion. Currently, the maximum possible flow rate of 460 µm s^−1^ is limited by the channel length and the time required for egress and invasion to occur, which is at the lower end of flow rates in the microvasculature. In the future, improved bonding between the glass and PDMS or a change in channel fabrication methodology could enable longer channels, thereby facilitating higher flow velocities and shear forces. The use of differential fluorescent labelling to distinguish multiple blood types or genetic variants could allow parallel testing, similar to the concept of grouping different blood types together in flow cytometry-based invasion assays [47]. This would enable systematic exploration of how different flow velocities affect the invasion of different red blood cell types, such as reticulocytes, compared with the mature erythrocytes used here or erythrocytes from specific blood groups or genetic backgrounds associated with protection against severe malaria. There is also considerable scope to modify the channel design to investigate the impact of a wide range of vessel geometries on invasion. This could include the introduction of junctions or constrictions, and channel diameter could be varied to mimic different vessel dimensions, especially given that this impacts the movement of the blood [48]. Sequestration of schizonts could also be mimicked by introducing pillar-cages into the channel. Late-stage infected erythrocytes are more rigid than uninfected erythrocytes [49, 50], which means that channel restrictions could be set such that erythrocytes, rings and merozoites could pass through, but schizonts could not. Finally, the automated invasion quantification presented here could be combined with microfluidic devices that more closely recapitulate the cellular environment, such as those containing epithelial cells that are currently being used to investigate sequestration [51, 52, 53].

## 4 Conclusion

In summary, we establish a microfluidic approach that enables the direct observation of malaria parasite invasion under flow, providing insight into how shear influences parasite–host interactions. Using this system, we show that loss of the invasion ligand PfEBA175 increases sensitivity to flow, likely due to changes in hydrodynamic forces opposing attachment rather than changes in contact frequency. In contrast, wild-type and ∆PfRH4 strains remain largely unaffected across the same flow regime. This shows that specific ligands may be particularly important for invasion under certain flow conditions in the bloodstream, a significance that cannot be captured by observing invasion in static assays alone.

## 5 Materials and Methods

### 5.1 Parasite culture

*P. falciparum* strains (either the wild-type NF54, 3D7 or genetically modified lines ∆PfRH4 or ∆PfEBA175 [36]) were cultured in human erythrocytes purchased from NHS Blood and Transplant, Cambridge, UK under the University of Cambridge Human Biology Research Ethics Committee, HBREC.2019.40 and ethical approval from NHS Cambridge South Research Ethics Committee, 20/EE/0100; the NHS obtained formal written consent for sample collection. The culture was kept at 37 ^*◦*^C in a gassed incubator or gassed and sealed culture container under a low oxygen atmosphere of 1% O_2_, 3% CO_2_, and 96% N_2_ (BOC, Guildford, UK). Cultures were routinely maintained at a 4% haematocrit in RPMI 1640 media (Gibco, UK) (routine culture was done using RPMI with phenol red and imaging was done without Phenol red which can cause fluorescent background) supplemented with 5 g L^−1^ Albumax II, DEXTROSE ANHYDROUS EP 2 g L^−1^, HEPES 5.96 g L^−1^, sodium bicarbonate EP 0.3 g L^−1^ and 0.05 g L^−1^ hypoxanthine dissolved in 2M NaOH.

### 5.2 Sorbitol synchronisation

Sorbitol synchronisation, described by [54], was performed by pelleting a culture with a large number of ring stage parasites, removing the supernatant and resuspending the pellet in ten times the pellet volume of 5% D-sorbitol (Sigma-Aldrich). This was incubated at 37 ^*◦*^C for 5 minutes, during which time later-stage parasites (trophozoites and schizonts) will rupture. The cells are then pelleted by centrifugation and then resuspended in RPMI at twenty times the pelleted volume. The cells were pelleted again and then resuspended in complete media to give 4% haematocrit.

### 5.3 Tight synchronisation of parasitised erythrocytes

After the lines were defrosted, they were sorbitol synchronised in the first cycle when the culture was greater than 0.5% rings and then delayed at room temperature to push the egress window into the intended synchronisation cycle. The parasites were tightly synchronised to a 3-hour window; before beginning, smears were checked to confirm the culture was predominantly schizonts with few rings. The media was removed from the culture, leaving 5 mL into which the infected blood was resuspended. This is then layered over 5 mL of 70% Percoll in a 15 mL tube (Percoll Merck P4937, 10% 10x PBS, 20% RPMI) and centrifuged at 1450 rcf for 11 minutes with brake set to 1 and acceleration to 3. This results in a band of late-stage parasites at the Percoll-media interface, which is removed and added to a flask with media and blood at a 4% haematocrit and incubated in a gassed incubator at 37 ^*◦*^C for 3 hours. The Percoll separation was then repeated as above, but this time, the bottom pellet containing the newly invaded ring stage parasites was kept. Then, sorbitol synchronisation was carried out to remove any remaining late-stage parasites that had not been removed by Percoll separation. This results in parasites that are all within a 3-hour developmental window, as they must have invaded new erythrocytes within the 3 hours post Percoll-purification. The parasites were then resuspended in 80 mL culture at 2.5% haematocrit. Once the parasitemia was higher than 2 % half of the parasites were kept from each cycle to reinvade, and half were used for microfluidics. The parasites for microfluidics were delayed for 28 hours at room temperature so that the imaging could be done on a different day from the synchronisation.

### 5.4 Printing of photoresist on silicon wafer

The mould used to cast the channels was created using soft lithography [55]. The positive of the design was first printed on photoresist film on a silicon wafer. A 11 mm Silicon Wafer (P(100) 0-100 Ω cm 500 µm SSP, University wafer Inc) was first washed with acetone and Isopropanol and then blow-dried with a Nitrogen gun. The wafer was then placed on a hot plate for 10 minutes at 200 ^*◦*^C before being left to cool on a lint-free wipe for 90 s at room temperature. The wafer was placed on the chuck of the spin coater (Laurell Technologies Corporation, model WS-650MZ-23NPP8) and 3 mL of SU8 6005 (A-Gas) was placed into the centre of the wafer and then spun for 10 s, at 500 rpm acceleration to 100 rpm/s, then for 30 s at 1900 rpm acceleration to 500 rpm/s. The coating was inspected to ensure an even application, and the process was repeated if any defects were observed. It was then soft-baked on a hot plate at 65 ^*◦*^C for 1 min; then the hot plate was turned up to 110 ^*◦*^C for a further 6 min.

The pattern of the channel was created using a Durham Magneto Optics Microwriter ML 3 to cross-link the photoresist in the area of the channels. A 50 µm resolution was used. Following printing, the wafer was post-exposure baked on a hot plate at 65 ^*◦*^C for 1 min, 95 ^*◦*^C for 6 min, then 65 ^*◦*^C for 1 min. Next, the unexposed photoresist was washed away. It was placed in the SU8 developer (PGMEA (Propylene glycol methyl ether acetate)) for 210 s or until the residue was completely removed and then washed in isopropanol. The wafer was hard-baked on a hot plate at 200 ^*◦*^C for 5 minutes. The depth of the channel was then measured using a profilometer (DektaXT, Bruker), resulting in a channel thickness of 6-7 µm. The shallow channels ensure a flat layer of erythrocytes, which facilitates their automated tracking.

### 5.5 Casting Polydimethylsiloxane (PDMS) and assembling devices

The silicon wafer was placed in a square plastic petri dish and visually inspected to check that it was completely free of dust. Polydimethylsiloxane (PDMS) (Sylgard 184 silicone elastomer kit) was made up by vigorously mixing the base with the curing agent (10% of the volume of base). It was degassed in a vacuum chamber and poured over the wafer. The PDMS was then cured at 60 ^*◦*^C for 2 hours. As multiple devices are on a single wafer, the individual devices are next cut out with a scalpel with roughly 0.5 cm of PDMS around the channels. The inlet and outlet holes were then punched using a 0.75 mm hole (Syneo CR0350255N20R4) and a Syneo punch machine. Debris blocking the narrow channels was a major issue, fig. S3, a large source of this was punching the holes; we found the debris was reduced by cutting holes over a clean piece of PDMS and washing the devices thoroughly after holes were punched. The PDMS devices and glass slides (Academy glass coverslips, 24 x 32 mm, 0.13-0.16 mm thick, 400-03-19) were washed with acetone, isopropanol and then deionised water, then dried using a nitrogen gun and placed channel side up in a petri dish in a 60 ^*◦*^C oven for 10 minutes. The PDMS device is then bonded to the glass slide using plasma treatment. The clean glass slide and the PDMS device with the channel side up are placed on the tray of the plasma cleaner (Harrick Plasma, Plasma Cleaner PDC-32G and Vacuum Gauge PDC-VCG-2). The chamber was depressurised to 0.1 with the valve down, and then the pressure was set to 0.5 with a valve to the right. The plasma was then activated on high for 30 s. After bonding, the device was placed for 10 min on a hotplate set to 90 ^*◦*^C and then left overnight in an oven at 60 ^*◦*^C. This step of heating the device after bonding was the factor that most significantly improved our bonding.

### 5.6 Setup of microfluidics device

Inlet and outlet tubes are connected to the holes in the PDMS devices using PDMS couplers (20G stainless steel 90 ^*◦*^C bent, Darwin Microfluidics PN-BEN-20G-100) and Tygon-tubing (ID 0.020 in OD 0.060 in, Cole-Palmer ND-100-80). Clamps were added to the two inlet tubes. Loading was done with a luer lock syringe needle (23G blunt-end, Darwin Microfluidics AE-23G-100x). With the pump inlet closed, the channel is blocked by loading 7.5% Bovine Serum Albumin (BSA) in Phosphate-buffered saline (PBS) (Merck A8412) into the loading inlet, and the device was placed at 37 ^*◦*^C for 30 min. Complete Phenol-Red free media was then passed through the channel.

### 5.7 Microscopy

A field of view close to the end of the channels was imaged at 37 ^*◦*^C using a custom-built, open-frame inverted microscope for 40 minutes. The objective used was a 20x air objective (Plan Apo VC 20x/ 0.75 DIC N2 OFN25 WD 1.0). Videos were recorded with alternate frames in brightfield (528 nm) and fluorescence (470 nm) illumination at 100 frames per second, using an LED bandpass fluorescent filter cube. The camera used was a Teledyne FLIR BFS-U3-70S7M-C with a 7.1 MP Sony IMX428 monochrome image sensor, resulting in an effective resolution of 0.224 µm*/*pixel. All images were captured at 3208 *×* 1000px, utilising only a portion of the sensor’s rows to achieve higher frame rates.

The flow in the channel is driven by a pressure pump (Disc pump evaluation kit P1329 020 XPSZ8 TTP Ventus/The Lee Company). The pressure pump was set to generate 19.2 kPa at the inlet. The outlet tubes were connected to columns of water of different heights to generate the backpressure that leads to different flow velocities in the channels. The column heights used were 0, 50, 100 and 150 cm, with the highest water column corresponding to 14.7 kPa added backpressure and thus the slowest flow velocity. The columns were placed on three-way valves, allowing the removal of the weight of the water columns when loading the microfluidic device with its malaria sample, so that all the channels could be loaded at the same rate.

### 5.8 Preparation of sample for imaging

The steps for setting up the sample for imaging are summarised in fig. S4. The tightly synchronised schizonts were arrested prior to egress by first isolating late-stage parasites from the culture using Percoll isolation as described above and then resuspending them in 20 mL of complete media with 1 µmol Compound 2 [30]. The samples were kept at 37 ^*◦*^C for a minimum of 3 hours and a maximum of 5.5 hours. This incubation window ensures that the majority of the parasites are paused in development just before egress. For each round of imaging, 10 mL of culture was removed from the flask, pelleted by centrifugation, and 9 mL of media was removed. The pellet was then resuspended in the remaining 1 mL of the media, ensuring that Compound 2 remained present. Next, SYBR Green (Invitrogen, Paisley, UK) was added to give a 1-in-5000 dilution and then incubation was carried out at 37 ^*◦*^C for 30 min. Two DNA dyes were tested, SYBR Green and Hoechst 33342 and it was shown that the SYBR Green gave a much brighter signal, fig. S2. Blood was added to give a 10% or 15% haematocrit, and the parasites were washed twice in 1 mL of complete Phenol-Red free media to remove Compound 2. Once Compound 2 is removed, merozoites start egressing after roughly 15 min. The culture was loaded into the channel as quickly as possible, with washing and loading usually taking about 10 min. The clamp on the input tubing was then closed, and the clip connected to the pressure pump was opened, with the pump already started. The valves for the outlet tubing were then set so that the water columns were connected to the outlet tubes. The pressure pump was then started, and once it was confirmed that the cells were flowing as expected, the camera recording was started and continued for at least 30 min to allow for invasion to occur.

### 5.9 Image analysis

The videos of the cells passing through the microfluidic channels were analysed using automated image analysis to identify the different cell types, and their trajectory across frames was tracked.

#### Finding erythrocytes

The first step is to analyse the brightfield images to detect all erythrocytes. The frames are pre-processed with a Gaussian blur. Then, we find the erythrocytes using the Hough circles implementation in OpenCV, [56] which uses gradient information from edges in the image. The parameters used are: minimum separation = 12 px, parameter 1 = 300, parameter 2 = 12, min radius = 8 px, and max radius = 30 px. This approach is effective, as we know the erythrocytes have a round shape, and looking for the contour enables the segmentation of cells that overlap.

#### Finding schizonts, merozoites, and rings

The fluorescent frames are next used to detect the parasites (schizonts, merozoites, and rings) as they are all stained with the SYBR Green DNA dye. This segmentation was performed using edge detection. First, the frames are pre-processed with a Gaussian blur and normalised so each pixel gets values between 0 and 255. A binary threshold determines all pixels with intensity above 28 as foreground, and a second Gaussian blur is applied to filter out spurious pixels and smooth the outline of true fluorescent signals. In order to filter out any background fluorescence, the contours are then detected and filtered to keep only elements with an area greater than 4 px.

#### Distinguishing schizonts and cell debris from merozoites and rings

Schizonts contain 26–32 merozoites within a single erythrocyte[3, 2], resulting in a significantly higher DNA content compared to a single merozoite or ring. Consequently, schizonts produce a much brighter fluorescent signal than merozoites or rings, both in terms of intensity and size.

Cells were classified as merozoites if their intensity per area fell below all defined linear boundaries for the merozoite population. Conversely, cells were categorised as schizonts if their intensity per area exceeded all defined linear boundaries for the schizont population. Any cells that did not satisfy the criteria for either category were classified as debris. The boundaries were manually selected to optimise the separation of rings and schizonts from debris, based on human-labelled datasets, fig. S5.

#### Distinguishing merozoites from rings

Distinguishing free merozoites from rings is challenging due to their similar fluorescent profiles, making classification based solely on fluorescent images unreliable. To address this, we leveraged the fact that the fluorescent signal of a ring is located within an erythrocyte. However, this approach is complicated by the frequent movement of merozoites near, above, or below erythrocytes within the channels, which can lead to misclassification. To overcome this, we used Trackpy’s NearestVelocityPredict [57] feature to link segmented masks into cell trajectories. For classification, any erythrocyte and merozoite that remained within their combined radius for the entire duration of their tracked trajectories (and for at least 40 ms) were considered “matched” and reclassified as rings.

### 5.10 Data availability

All code and raw data supporting the findings of this study are available in the Zenodo repository at https://doi.org/10.5281/zenodo.15399769.

## Acknowledgements

We would like to thank Theresa Feltwell, Nadia Cross and the CIMR and Physics of Medicine Department support staff for their invaluable logistical support; Yen Chun Lin and Margherita Tavasso who worked on previous versions of microfluidic devices to follow malaria invasion which provided important proofs of concept; Dean Kos and Ishaan Lohia for their assistance and training whilst they were the microfluidics facility managers. This work was funded by Wellcome, grant numbers (220266/Z/20/Z to JR), (222323/Z/21/Z to EK) and (20485/Z/16/Z to VI); EU-EC Marie Curie ITN PyMot (955910 to MK) and Engineering and Physical Sciences Research Council (EPSRC) grant (EP/W004453/1 to PC). For the purpose of open access, the author has applied a CC BY copyright license to any Author Accepted Manuscript Version arising from this submission. The funders played no role in the study design, data collection and analysis, decision to publish, or preparation of the manuscript.

## References

[1] WHO. World malaria report 2022. WHO, 2022. ISBN 9789240040496. URL https://www.who.int/teams/global-malaria-programme/reports/world-malaria-report-2021.

[2] Svetlana Glushakova, Amanda Balaban, Philip G. McQueen, Rosane Coutinho, Jeffery L. Miller, Ralph Nossal, Rick M. Fairhurst, and Joshua Zimmerberg. Hemoglobinopathic erythrocytes affect the intraerythrocytic multiplication of Plasmodium falciparum in vitro. Journal of Infectious Diseases, 210(7):1100–1109, 2014. ISSN 15376613. doi: 10.1093/infdis/jiu203.

[3] Rachel M. Rudlaff, Stephan Kraemer, Jeffrey Marshman, and Jeffrey D. Dvorin. Three-dimensional ultrastructure of Plasmodium falciparum throughout cytokinesis. PLoS Pathogens, 16(6):1–21, 2020. ISSN 15537374. doi: 10.1371/journal.ppat.1008587. URL http://dx.doi.org/10.1371/journal.ppat.1008587.

[4] Paul R. Gilson and Brendan S. Crabb. Morphology and kinetics of the three distinct phases of red blood cell invasion by Plasmodium falciparum merozoites. International Journal for Parasitology, 39(1):91–96, 2009. ISSN 00207519. doi: 10.1016/j.ijpara.2008.09.007. URL http://dx.doi.org/10.1016/j.ijpara.2008.09.007.

[5] M. Aikawa, M. Iseki, J. W. Barnwell, D. Taylor, M. M. Oo, and R. J. Howard. The pathology of human cerebral malaria. American Journal of Tropical Medicine and Hygiene, 43(2 SUPPL.):30–37, 1990. ISSN 00029637. doi: 10.4269/ajtmh.1990.43.30.

[6] Regina Joice, Sandra K Nilsson, Jacqui Montgomery, Selasi Dankwa, Belinda Morahan, Karl B Seydel, Lucia Bertuccini, Pietro Alano, C Kim, Manoj T Duraisingh, Terrie E Taylor, and Danny A Milner. Levels and changes of HDL cholesterol and apolipoprotein A-I in relation to risk of cardiovascular events among statin-treated patients; a meta-analysis. Circulation, 6(244):1–16, 2014. ISSN 15378276. doi: 10.1126/scitranslmed.3008882.Plasmodium.

[7] Oranan Prommano, Urai Chaisri, Gareth D H Turner, Polrat Wilairatana, David J P Ferguson, Parnpen Viriyavejakul, Nicholas J. White, and Emsri Pongponratn. A quantitative ultrastructural study of the liver and the spleen in fatal falciparum malaria. The Southeast Asian journal of tropical medicine and public health, 36(6):1359–70, 11 2005. ISSN 0125-1562. URL http://www.ncbi.nlm.nih.gov/pubmed/16610635.

[8] Britta C. Urban, Tran T. Hien, Nicholas P. Day, Nguyen H. Phu, Rachel Roberts, Emsri Pongponratn, Margret Jones, Nguyen T.H. Mai, Delia Bethell, Gareth D.H. Turner, David Ferguson, Nicholas J. White, and David J. Roberts. Fatal Plasmodium falciparum malaria causes specific patterns of splenic architectural disorganization. Infection and Immunity, 73(4):1986–1994, 2005. ISSN 00199567. doi: 10.1128/IAI.73.4.1986-1994.2005.

[9] Steven Kho, Labibah Qotrunnada, Leo Leonardo, Benediktus Andries, Putu A.I. Wardani, Aurelie Fricot, Benoit Henry, David Hardy, Nur I. Margyaningsih, Dwi Apriyanti, Agatha M. Puspitasari, Pak Prayoga, Leily Trianty, Enny Kenangalem, Fabrice Chretien, Innocent Safeukui, Hernando A. del Portillo, Carmen Fernandez-Becerra, Elamaran Meibalan, Matthias Marti, Ric N. Price, Tonia Woodberry, Papa A. Ndour, Bruce M. Russell, Tsin W. Yeo, Gabriela Minigo, Rintis Noviyanti, Jeanne R. Poespoprodjo, Nurjati C. Siregar, Pierre A. Buffet, and Nicholas M. Anstey. Hidden Biomass of Intact Malaria Parasites in the Human Spleen. New England Journal of Medicine, 384(21): 2067–2069, 5 2021. ISSN 0028-4793. doi: 10.1056/NEJMc2023884. URL http://www.nejm.org/doi/10.1056/NEJMc2023884.

[10] Aristotle G. Koutsiaris, Sophia V. Tachmitzi, and Nick Batis. Wall shear stress quantification in the human conjunctival pre-capillary arterioles in vivo. Microvascular Research, 85(1):34–39, 2013. ISSN 00262862. doi: 10.1016/j.mvr.2012.11.003. URL http://dx.doi.org/10.1016/j.mvr.2012.11.003.

[11] Aristotle G. Koutsiaris, Sophia V. Tachmitzi, Nick Batis, Maria G. Kotoula, Constantinos H. Karabatsas, Evagelia Tsironi, and Dimitrios Z. Chatzoulis. Volume flow and wall shear stress quantification in the human conjunctival capillaries and post-capillary venules in vivo. Biorheology, 44(5-6):375–86, 2007. ISSN 0006-355X. URL http://www.ncbi.nlm.nih.gov/pubmed/18401076.

[12] M. Klarhöfer, B. Csapo, Cs Balassy, J. C. Szeles, and E. Moser. High-resolution blood flow velocity measurements in the human finger. Magnetic Resonance in Medicine, 45(4):716–719, 2001. ISSN 07403194. doi: 10.1002/mrm.1096.

[13] G.A. Butcher. The behaviour of different strains of Plasmodium falciparum in suspension and static culture. Transactions of the Royal Society of Tropical Medicine and Hygiene, 76(3):407–409, 1 1982. ISSN 00359203. doi: 10.1016/0035-9203(82)90202-4. URL https://academic.oup.com/trstmh/article-lookup/doi/10.1016/0035-9203(82)90202-4.

[14] J. Werner Zolg, Alexander J. MacLeod, Ida H. Dickson, and John G. Scaife. Plasmodium falciparum: Modifications of the In vitro Culture Conditions Improving Parasitic Yields. The Journal of Parasitology, 68(6):1072, 12 1982. ISSN 00223395. doi: 10.2307/3281094. URL https://www.jstor.org/stable/3281094?origin=crossref.

[15] Richard J.W. Allen and Kiaran Kirk. Plasmodium falciparum culture: The benefits of shaking. Molecular and Biochemical Parasitology, 169(1):63–65, 1 2010. ISSN 01666851. doi: 10.1016/j.molbiopara.2009.09.005. URL https://linkinghub.elsevier.com/retrieve/pii/S0166685109002205.

[16] Ulf Ribacke, Kirsten Moll, Letusa Albrecht, Hodan Ahmed Ismail, Johan Normark, Emilie Flaberg, Laszlo Szekely, Kjell Hultenby, Kristina E. M. Persson, Thomas G. Egwang, and Mats Wahlgren. Improved In Vitro Culture of Plasmodium falciparum Permits Establishment of Clinical Isolates with Preserved Multiplication, Invasion and Rosetting Phenotypes. PLoS ONE, 8(7):e69781, 7 2013. ISSN 1932-6203. doi: 10.1371/journal.pone.0069781. URL https://linkinghub.elsevier.com/retrieve/pii/S1471492220302245 https://dx.plos.org/10.1371/journal.pone.0069781.

[17] Gordon A. Awandare, Prince B. Nyarko, Yaw Aniweh, Reuben Ayivor-Djanie, and José A. Stoute. Plasmodium falciparum strains spontaneously switch invasion phenotype in suspension culture. Scientific Reports, 8(1):5782, 4 2018. ISSN 2045-2322. doi: 10.1038/s41598-018-24218-0. URL https://www.nature.com/articles/s41598-018-24218-0.

[18] Barbara Clough, F. Abiola Atilola, and Geoffrey Pasvol. The role of rosetting in the multiplication of Plasmodium falciparum : rosette formation neither enhances nor targets parasite invasion into uninfected red cells. British Journal of Haematology, 100(1):99–104, 1 1998. ISSN 0007-1048. doi: 10.1046/j.1365-2141.1998.00534.x. URL https://onlinelibrary.wiley.com/doi/10.1046/j.1365-2141.1998.00534.x.

[19] Emma Kals, Morten Kals, Pietro Cicuta, and Julian C Rayner. Under conditions of high wall shear stress, several PfEBA and PfRH ligands are important for malaria Plasmodium falciparum blood-stage growth. mBio, 16(8), 8 2025. ISSN 2150-7511. doi: 10.1128/mbio.01499-25. URL https://journals.asm.org/doi/10.1128/mbio.01499-25.

[20] Rob Driessen, Feihu Zhao, Sandra Hofmann, Carlijn Bouten, Cecilia Sahlgren, and Oscar Stassen. Computational Characterization of the Dish-In-A-Dish, A High Yield Culture Platform for Endothelial Shear Stress Studies on the Orbital Shaker. Micromachines, 11(6):552, 5 2020. ISSN 2072-666X. doi: 10.3390/mi11060552. URL https://www.mdpi.com/2072-666X/11/6/552.

[21] Bernhard Sebastian and Petra S. Dittrich. Microfluidics to Mimic Blood Flow in Health and Disease. Annual Review of Fluid Mechanics, 50(1):483–504, 1 2018. ISSN 0066-4189. doi: 10.1146/annurev-fluid-010816-060246. URL https://www.annualreviews.org/doi/10.1146/annurev-fluid-010816-060246.

[22] J. Patrick Shelby, John White, Karthikeyan Ganesan, Pradipsinh K. Rathod, and Daniel T. Chiu. A microfluidic model for single-cell capillary obstruction by Plasmodium falciparum-infected erythrocytes. Proceedings of the National Academy of Sciences, 100(25):14618–14622, 12 2003. ISSN 0027-8424. doi: 10.1073/pnas.2433968100. URL https://pnas.org/doi/full/10.1073/pnas.2433968100.

[23] Thurston Herricks, Meher Antia, and Pradipsinh K. Rathod. Deformability limits of Plasmodium falciparum-infected red blood cells. Cellular Microbiology, 11(9):1340–1353, 2009. ISSN 14625814. doi: 10.1111/j.1462-5822.2009.01334.x.

[24] Quan Guo, Sarah J. Reiling, Petra Rohrbach, and Hongshen Ma. Microfluidic biomechanical assay for red blood cells parasitized by Plasmodium falciparum. Lab on a Chip, 12(6):1143–1150, 3 2012. ISSN 14730189. doi: 10.1039/c2lc20857a.

[25] Majid Ebrahimi Warkiani, Andy Kah Ping Tay, Bee Luan Khoo, Xu Xiaofeng, Jongyoon Han, and Chwee Teck Lim. Malaria detection using inertial microfluidics. Lab on a Chip, 15(4):1101–1109, 2015. ISSN 14730189. doi: 10.1039/c4lc01058b.

[26] T. M. Geislinger, B. Eggart, S. Braunmüller, L. Schmid, and T. Franke. Separation of blood cells using hydrodynamic lift. Applied Physics Letters, 100(18), 2012. ISSN 00036951. doi: 10.1063/1.4709614.

[27] Ying Bena Lim, Juzar Thingna, Jianshu Cao, and Chwee Teck Lim. Single molecule and multiple bond characterization of catch bond associated cytoadhesion in malaria. Scientific Reports, 7(1):1–11, 2017. ISSN 20452322. doi: 10.1038/s41598-017-04352-x.

[28] Meher Antia, Thurston Herricks, and Pradipsinh K. Rathod. Microfluidic modeling of cell-cell interactions in malaria pathogenesis. PLoS Pathogens, 3(7):0939–0948, 2007. ISSN 15537374. doi: 10.1371/journal.ppat.0030099.

[29] Maria Bernabeu, Celina Gunnarsson, Maria Vishnyakova, Caitlin C. Howard, Ryan J. Nagao, Marion Avril, Terrie E Taylor, Karl B Seydel, Ying Zheng, and Joseph D. Smith. Binding Heterogeneity of Plasmodium falciparum to Engineered 3D Brain Microvessels Is Mediated by EPCR and ICAM-1. mBio, 10(3):1–16, 6 2019. ISSN 2161-2129. doi: 10.1128/mBio.00420-19. URL https://journals.asm.org/doi/10.1128/mBio.00420-19.

[30] Margarida Ressurreição, James A. Thomas, Stephanie D. Nofal, Christian Flueck, Robert W. Moon, David A. Baker, and Christiaan van Ooij. Use of a highly specific kinase inhibitor for rapid, simple and precise synchronization of Plasmodium falciparum and Plasmodium knowlesi asexual blood-stage parasites. PLOS ONE, 15(7):e0235798, 7 2020. ISSN 1932-6203. doi: 10.1371/journal.pone.0235798. URL https://dx.plos.org/10.1371/journal.pone.0235798.

[31] David Walliker, Isabella A. Quakyi, Thomas E. Wellems, Thomas F. McCutchan, Ana Szarfman, William T. London, Lynn M. Corcoran, Thomas R. Burkot, and Richard Carter. Genetic Analysis of the Human Malaria Parasite Plasmodium falciparum. Science, 236(4809):1661–1666, 6 1987. ISSN 0036-8075. doi: 10.1126/science.3299700. URL http://www.sciencemag.org/cgi/doi/10.1126/science.3299700 https://www.science.org/doi/10.1126/science.3299700.

[32] Kara A. Moser, Elliott F. Drábek, Ankit Dwivedi, Emily M. Stucke, Jonathan Crabtree, Antoine Dara, Zalak Shah, Matthew Adams, Tao Li, Priscila T. Rodrigues, Sergey Koren, Adam M. Phillippy, James B. Munro, Amed Ouattara, Benjamin C. Sparklin, Julie C. Dunning Hotopp, Kirsten E. Lyke, Lisa Sadzewicz, Luke J. Tallon, Michele D. Spring, Krisada Jongsakul, Chanthap Lon, David L. Saunders, Marcelo U. Ferreira, Myaing M. Nyunt, Miriam K. Laufer, Mark A. Travassos, Robert W. Sauerwein, Shannon Takala-Harrison, Claire M. Fraser, B. Kim Lee Sim, Stephen L. Hoffman, Christopher V. Plowe, and Joana C. Silva. Strains used in whole organism Plasmodium falciparum vaccine trials differ in genome structure, sequence, and immunogenic potential. Genome Medicine, 12(1):6, 12 2020. ISSN 1756-994X. doi: 10.1186/s13073-019-0708-9. URL https://genomemedicine.biomedcentral.com/articles/10.1186/s13073-019-0708-9.

[33] Catherine J. Merrick, Rays H. Y. Jiang, Kristen M. Skillman, Upeka Samarakoon, Rachel M. Moore, Ron Dzikowski, Michael T. Ferdig, and Manoj T. Duraisingh. Functional Analysis of Sirtuin Genes in Multiple Plasmodium falciparum Strains. PLOS ONE, 10(3):e0118865, 3 2015. ISSN 1932-6203. doi: 10.1371/journal.pone.0118865. URL https://dx.plos.org/10.1371/journal.pone.0118865.

[34] Michael J. Delves, Ursula Straschil, Andrea Ruecker, Celia Miguel-Blanco, Sara Marques, Alexandre C. Dufour, Jake Baum, and Robert E. Sinden. Routine in vitro culture of P. Falciparum gametocytes to evaluate novel transmission-blocking interventions. Nature Protocols, 11(9):1668–1680, 2016. ISSN 17502799. doi: 10.1038/nprot.2016.096.

[35] Sabine A. Fraschka, Michael Filarsky, Regina Hoo, Igor Niederwieser, Xue Yan Yam, Nicolas M.B. Brancucci, Franziska Mohring, Annals T. Mushunje, Ximei Huang, Peter R. Christensen, Francois Nosten, Zbynek Bozdech, Bruce Russell, Robert W. Moon, Matthias Marti, Peter R. Preiser, Richárd Bártfai, and Till S. Voss. Comparative Heterochromatin Profiling Reveals Conserved and Unique Epigenome Signatures Linked to Adaptation and Development of Malaria Parasites. Cell Host & Microbe, 23(3):407–420, 3 2018. ISSN 19313128. doi: 10.1016/j.chom.2018.01.008. URL https://linkinghub.elsevier.com/retrieve/pii/S193131281830043X.

[36] Emma Kals, Morten Kals, Rebecca A Lees, Viola Introini, Alison Kemp, Eleanor Silvester, Christine R Collins, Trishant Umrekar, Jurij Kotar, Pietro Cicuta, and Julian C Rayner. Application of optical tweezer technology reveals that PfEBA and PfRH ligands, not PfMSP1, play a central role in Plasmodium falciparum merozoite-erythrocyte attachment. PLOS Pathogens, 20(9):e1012041, 9 2024. ISSN 1553-7374. doi: 10.1371/journal.ppat.1012041. URL http://biorxiv.org/content/early/2024/02/13/2024.02.13.580055.abstract https://dx.plos.org/10.1371/journal.ppat.1012041.

[37] Wai Hong Tham, Julie Healer, and Alan F. Cowman. Erythrocyte and reticulocyte binding-like proteins of Plasmodium falciparum. Trends in Parasitology, 28(1):23–30, 2012. ISSN 14714922. doi: 10.1016/j.pt.2011.10.002. URL http://dx.doi.org/10.1016/j.pt.2011.10.002.

[38] Sash Lopaticki, Alexander G. Maier, Jennifer Thompson, Danny W. Wilson, Wai-Hong Tham, Tony Triglia, Alex Gout, Terence P. Speed, James G. Beeson, Julie Healer, and Alan F. Cowman. Reticulocyte and Erythrocyte Binding-Like Proteins Function Cooperatively in Invasion of Human Erythrocytes by Malaria Parasites. Infection and Immunity, 79(3):1107–1117, 3 2011. ISSN 0019-9567. doi: 10.1128/IAI.01021-10. URL https://journals.asm.org/doi/10.1128/IAI.01021-10.

[39] Kristina E.M. Persson, Fiona J. McCallum, Linda Reiling, Nicole A. Lister, Janine Stubbs, Alan F. Cowman, Kevin Marsh, and James G. Beeson. Variation in use of erythrocyte invasion pathways by Plasmodium falciparum mediates evasion of human inhibitory antibodies. Journal of Clinical Investigation, 118(1):342–351, 1 2008. ISSN 0021-9738. doi: 10.1172/JCI32138. URL http://content.the-jci.org/articles/view/32138.

[40] Alexander G. Maier, Manoj T. Duraisingh, John C. Reeder, Sheral S. Patel, James W. Kazura, Peter A. Zimmerman, and Alan F. Cowman. Plasmodium falciparum erythrocyte invasion through glycophorin C and selection for Gerbich negativity in human populations. Nature Medicine, 9(1):87–92, 2003. ISSN 10788956. doi: 10.1038/nm807.

[41] Tony Triglia, Manoj T. Duraisingh, Robert T. Good, and Alan F. Cowman. Reticulocyte-binding protein homologue 1 is required for sialic acid-dependent invasion into human erythrocytes by Plasmodium falciparum. Molecular Microbiology, 55(1):162–174, 1 2005. ISSN 0950-382X. doi: 10.1111/j.1365-2958.2004.04388.x. URL https://onlinelibrary.wiley.com/doi/10.1111/j.1365-2958.2004.04388.x.

[42] Manoj T. Duraisingh, Tony Triglia, Stuart A. Ralph, Julian C. Rayner, John W. Barnwell, Geoffrey I. McFadden, and Alan F. Cowman. Phenotypic variation of Plasmodium falciparum merozoite proteins directs receptor targeting for invasion of human erythrocytes. The EMBO Journal, 22(5):1047–1057, 3 2003. ISSN 0261-4189. doi: 10.1093/emboj/cdg096. URL http://emboj.embopress.org/cgi/doi/10.1093/emboj/cdg096.

[43] Francis W Klotz, Palmer A Orlandi, Gerd Reuter, Stuart J Cohen, J.David Haynes, Roland Schauer, Russell J Howard, Peter Palese, and Louis H Miller. Binding of Plasmodium falciparum 175-kilodalton erythrocyte binding antigen and invasion of murine erythrocytes requires N-acetylneuraminic acid but not its O-acetylated form. Molecular and Biochemical Parasitology, 51(1):49–54, 3 1992. ISSN 01666851. doi: 10.1016/0166-6851(92)90199-T. URL https://linkinghub.elsevier.com/retrieve/pii/016668519290199T.

[44] B. K.L. Sim, C. E. Chitnis, K. Wasniowska, T. J. Hadley, and L. H. Miller. Receptor and ligand domains for invasion of erythrocytes by Plasmodium falciparum. Science, 264(5167):1941–1944, 1994. ISSN 00368075. doi: 10.1126/science.8009226.

[45] Ewa Jaskiewicz, Marlena Jodłowska, Radosław Kaczmarek, and Agata Zerka. Erythrocyte glycophorins as receptors for Plasmodium merozoites. Parasites and Vectors, 12(1):1–11, 2019. ISSN 17563305. doi: 10.1186/s13071-019-3575-8. URL https://doi.org/10.1186/s13071-019-3575-8.

[46] Olsson Reid, Lomas-Francis. The blood group antigen factsbook. Amsterdam; Boston : Elsevier/Academic Press, 2012. ISBN 0-240-82130-0.

[47] Michel Theron, Richard L. Hesketh, Sathish Subramanian, and Julian C. Rayner. An adaptable two-color flow cytometric assay to quantitate the invasion of erythrocytes by Plasmodium falciparum parasites. Cytometry Part A, 77(11):1067–1074, 2010. ISSN 15524922. doi: 10.1002/cyto.a.20972.

[48] Timothy W. Secomb. Blood Flow in the Microcirculation. Annual Review of Fluid Mechanics, 49(1):443–461, 1 2017. ISSN 0066-4189. doi: 10.1146/annurev-fluid-010816-060302. URL https://www.annualreviews.org/doi/10.1146/annurev-fluid-010816-060302.

[49] GB B Nash, E O’brien, EC C Gordon-Smith, and JA A Dormandy. Abnormalities in the mechanical properties of red blood cells caused by Plasmodium falciparum. Blood, 74(2):855–861, 8 1989. ISSN 0006-4971. doi: 10.1182/blood.V74.2.855.855. URL http://dx.doi.org/10.1182/blood.V74.2.855.855 https://ashpublications.org/blood/article/74/2/855/167233/Abnormalities-in-the-mechanical-properties-of-red.

[50] Y. Park, Monica Diez-Silva, Gabriel Popescu, George Lykotrafitis, Wonshik Choi, Michael S. Feld, and Subra Suresh. Refractive index maps and membrane dynamics of human red blood cells parasitized by Plasmodium falciparum. Proceedings of the National Academy of Sciences, 105(37):13730–13735, 9 2008. ISSN 0027-8424. doi: 10.1073/pnas.0806100105. URL http://www.pnas.org/cgi/doi/10.1073/pnas.0806100105.

[51] Ying Zheng, Junmei Chen, and José A. López. Flow-driven assembly of VWF fibres and webs in in vitro microvessels. Nature Communications, 6, 2015. ISSN 20411723. doi: 10.1038/ncomms8858.

[52] Ying Zheng, Junmei Chen, Michael Craven, Nak Won Choi, Samuel Totorica, Anthony Diaz-Santana, Pouneh Kermani, Barbara Hempstead, Claudia Fischbach-Teschl, José A. López, and Abraham D. Stroock. In vitro microvessels for the study of angiogenesis and thrombosis. Proceedings of the National Academy of Sciences of the United States of America, 109(24):9342–9347, 2012. ISSN 00278424. doi: 10.1073/pnas.1201240109.

[53] Christopher Arakawa, Celina Gunnarsson, Caitlin Howard, Maria Bernabeu, Kiet Phong, Eric Yang, Cole A. DeForest, Joseph D. Smith, and Ying Zheng. Biophysical and biomolecular interactions of malaria-infected erythrocytes in engineered human capillaries. Science Advances, 6(3), 2020. ISSN 23752548. doi: 10.1126/sciadv.aay7243.

[54] Chris Lambros and Jerome P. Vanderberg. Synchronization of Plasmodium falciparum Erythrocytic Stages in Culture. The Journal of Parasitology, 65(3):418, 6 1979. ISSN 00223395. doi: 10.2307/3280287. URL https://www.jstor.org/stable/3280287?origin=crossref.

[55] Xiao-Mei Zhao, Younan Xia, and George M. Whitesides. Soft lithographic methods for nano-fabrication. Journal of Materials Chemistry, 7(7):1069–1074, 1997. ISSN 09599428. doi: 10.1039/a700145b. URL https://xlink.rsc.org/?DOI=a700145b.

[56] OpenCV. No Title, 2024. URL https://docs.opencv.org/4.x/da/d53/tutorial_py_houghcircles.html.

[57] Daniel Allan. soft-matter/trackpy: v0.6.4. 2024. doi: 10.5281/zenodo.1213240. URL https://doi.org/10.5281/zenodo.12708864.

