## Supplementary material for "Microfluidic Platform for Automatic Quantification of Malaria Parasite Invasion Under Physiological Flow Conditions": Suplemental information

#### This PDF file includes:

Supporting text
Figs. S1 to S10
Table S1

### Supporting Information Text

#### Section S1. Estimation of Hydrodynamic Forces on a Merozoite in the Microfluidic Channel

We estimate the characteristic hydrodynamic forces acting on a $\sim 1{-}2\ \mu$m merozoite as it encounters a red blood cell (RBC) within our microfluidic device. The channel dimensions are 6 $\mu$m in height and 100 $\mu$m in width, and the suspension (4% hematocrit in culture media) flows at an average velocity of $U_{\text{avg}} = 410\,\mu\text{m}\,\text{s}^{-1}$. The effective viscosity of the suspension is taken as $\mu \approx 1.2 \times 10^{-3}$ Pa · s, close to plasma viscosity at low hematocrit.

**Reynolds Number and Flow Regime.** Using the channel height $h = 6\ \mu$m as the characteristic length, the Reynolds number is

$$\text{Re} = \frac{\rho U_{\text{avg}} h}{\mu} \approx \frac{10^3 \times 4.5 \times 10^{-4} \times 6 \times 10^{-6}}{1.2 \times 10^{-3}} \approx 2 \times 10^{-3} \ll 1.$$

Thus, the flow is well within the creeping flow regime.

**Wall Shear Rate.** Because the channel is much wider than it is tall ($100\ \mu\text{m} \gg 6\ \mu\text{m}$), the flow can be approximated as plane Poiseuille flow between parallel plates. The wall shear rate for such a geometry is

$$\dot{\gamma}_{\text{wall}} \approx \frac{6\,U_{\text{avg}}}{h},$$

which yields

$$\dot{\gamma}_{\text{wall}} \approx \frac{6 \times 4.1 \times 10^{-4}}{6 \times 10^{-6}} \approx 410\ \text{s}^{-1}.$$

The corresponding wall shear stress is

$$\tau_{\text{wall}} = \mu \dot{\gamma}_{\text{wall}} \approx (1.2 \times 10^{-3})(410) \approx 0.5\ \text{Pa}.$$

**Hydrodynamic Force Estimates.** We model the merozoite as a particle of characteristic size $L \sim 1\ \mu$m. Two equivalent Stokes-flow–based approaches give similar force scales.

**(1) Shear Stress Acting over the Particle Area** Taking the exposed area as surface area of a sphere $A = \pi d^2 \approx 1.2 \times 10^{-11}\ \text{m}^2$,

$$F \approx \tau_{\text{wall}}\, A = 6.2 \times 10^{-12}\ \text{N} \approx 6\ \text{pN}$$

**(2) Stokes Drag in a Linear Shear Field** Near the wall, the velocity profile is approximately linear, giving a relative velocity $U_{\text{rel}} \sim \dot{\gamma}_{\text{wall}} L$. The Stokes drag on a sphere of radius $a \approx 1\ \mu$m in such a flow is

$$F \approx 6\pi\mu a U_{\text{rel}} = 6\pi\mu a^2 \dot{\gamma}_{\text{wall}}.$$

Substituting values,

$$F \approx 6\pi(1.2 \times 10^{-3})(10^{-6})^2(410) \approx 9.3 \times 10^{-12}\ \text{N} \approx 10\ \text{pN}.$$

**Resulting Force Scale.** Both methods produce consistent estimates around 10 pN.

**Table S1. Table showing error rates for each cell type, broken down for each frame and channel.**

| Strain | Repeat | Experiment | Channel | Time (sec) | Schizonts | Rings | Merozoites | Erythrocytes | Debris |
|---|---|---|---|---|---|---|---|---|---|
| KoRH4 NF54 | 3 | ECK17 | 0 | 25.19 | 0.50 | 0.00 | 0.00 | 0.02 | -1.00 |
| KoRH4 NF54 | 3 | ECK17 | 0 | 328.75 | - | 0.00 | -0.25 | 0.02 | -1.00 |
| KoRH4 NF54 | 3 | ECK17 | 1 | 25.19 | 0.00 | 0.00 | 0.00 | 0.12 | 0.00 |
| KoRH4 NF54 | 3 | ECK17 | 1 | 328.75 | -1.00 | -1.00 | 0.00 | 0.03 | - |
| KoRH4 NF54 | 3 | ECK17 | 2 | 25.19 | 0.00 | -1.00 | - | 0.17 | 0.00 |
| KoRH4 NF54 | 3 | ECK17 | 2 | 328.75 | 0.00 | 0.00 | 0.00 | 0.09 | 0.00 |
| KoRH4 NF54 | 3 | ECK17 | 3 | 25.19 | 0.00 | 0.00 | 0.00 | 0.02 | 0.00 |
| KoRH4 NF54 | 3 | ECK17 | 3 | 328.75 | 1.00 | 0.00 | 0.00 | 0.03 | -0.67 |
| NF54 | 4 | ECK25 | 0 | 25.43 | 2.00 | 0.00 | 0.00 | 0.14 | -1.00 |
| NF54 | 4 | ECK25 | 0 | 502.37 | 0.33 | -0.30 | 0.22 | -0.04 | -0.50 |
| NF54 | 4 | ECK25 | 1 | 25.43 | 0.00 | 0.00 | 0.00 | 0.25 | 0.00 |
| NF54 | 4 | ECK25 | 1 | 502.37 | - | -0.08 | -0.07 | 0.10 | -1.00 |
| NF54 | 4 | ECK25 | 2 | 25.43 | - | - | -0.50 | 0.26 | 0.00 |
| NF54 | 4 | ECK25 | 2 | 502.37 | - | -0.12 | 0.25 | 0.00 | -1.00 |
| NF54 | 4 | ECK25 | 3 | 25.43 | 2.00 | 0.00 | 0.00 | 0.26 | -1.00 |
| NF54 | 4 | ECK25 | 3 | 502.37 | 0.00 | -0.16 | 0.07 | 0.07 | 0.00 |
| 3D7 | 6 | ECK26 | 0 | 25.00 | 0.00 | 0.00 | 0.00 | 0.09 | -1.00 |
| 3D7 | 6 | ECK26 | 0 | 322.69 | - | 0.40 | -0.22 | 0.19 | -1.00 |
| 3D7 | 6 | ECK26 | 1 | 25.00 | -0.29 | 0.00 | - | -0.15 | -1.00 |
| 3D7 | 6 | ECK26 | 1 | 322.69 | 0.00 | 0.25 | 0.00 | 0.11 | 0.00 |
| 3D7 | 6 | ECK26 | 2 | 25.00 | - | 0.00 | 0.00 | 0.26 | 0.00 |
| 3D7 | 6 | ECK26 | 2 | 322.69 | - | 0.09 | 0.11 | 0.28 | -1.00 |
| 3D7 | 6 | ECK26 | 3 | 25.00 | - | 0.00 | -1.00 | 0.38 | -1.00 |
| 3D7 | 6 | ECK26 | 3 | 322.69 | 0.00 | 0.22 | -0.11 | 0.40 | 0.00 |
| NF54 | 5 | ECK27 | 0 | 51.00 | 0.00 | 0.00 | -0.50 | 0.18 | -1.00 |
| NF54 | 5 | ECK27 | 0 | 486.26 | 0.00 | -0.33 | 0.25 | 0.36 | 0.00 |
| NF54 | 5 | ECK27 | 1 | 51.00 | 1.00 | 0.00 | 0.00 | 0.27 | -1.00 |
| NF54 | 5 | ECK27 | 1 | 486.26 | 0.00 | -0.47 | 0.14 | 0.25 | 0.00 |
| NF54 | 5 | ECK27 | 2 | 51.00 | 2.00 | 0.00 | 0.00 | 0.49 | -1.00 |
| NF54 | 5 | ECK27 | 2 | 486.26 | - | -0.41 | 0.14 | 0.15 | 0.00 |
| NF54 | 5 | ECK27 | 3 | 51.00 | 0.00 | -1.00 | - | -0.03 | 0.00 |
| NF54 | 5 | ECK27 | 3 | 486.26 | - | -0.31 | 0.25 | 0.28 | - |

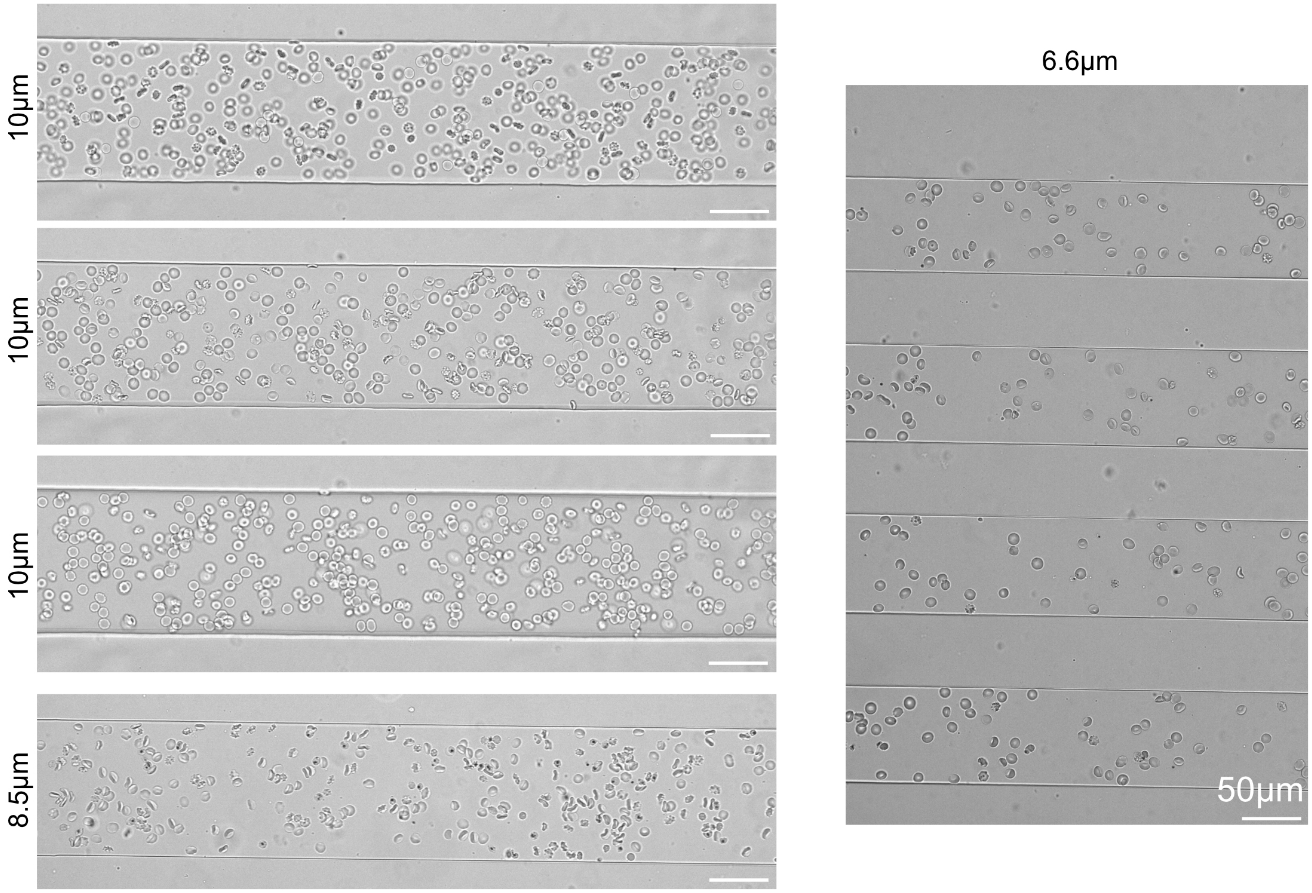


**Fig. S1. Examples of parasites imaged in channels of different depths.** Three images are shown for the 10 $\mu$m channel at three different depths of focus. The 8.5 $\mu$m and 6.6 $\mu$m channels are shown at a single focus. Error bars are 50 $\mu$m. The 10 $\mu$m and 8.5 $\mu$m deep channels show erythrocytes in the single-channel device with a 100 $\mu$m width. The 6.6 $\mu$m

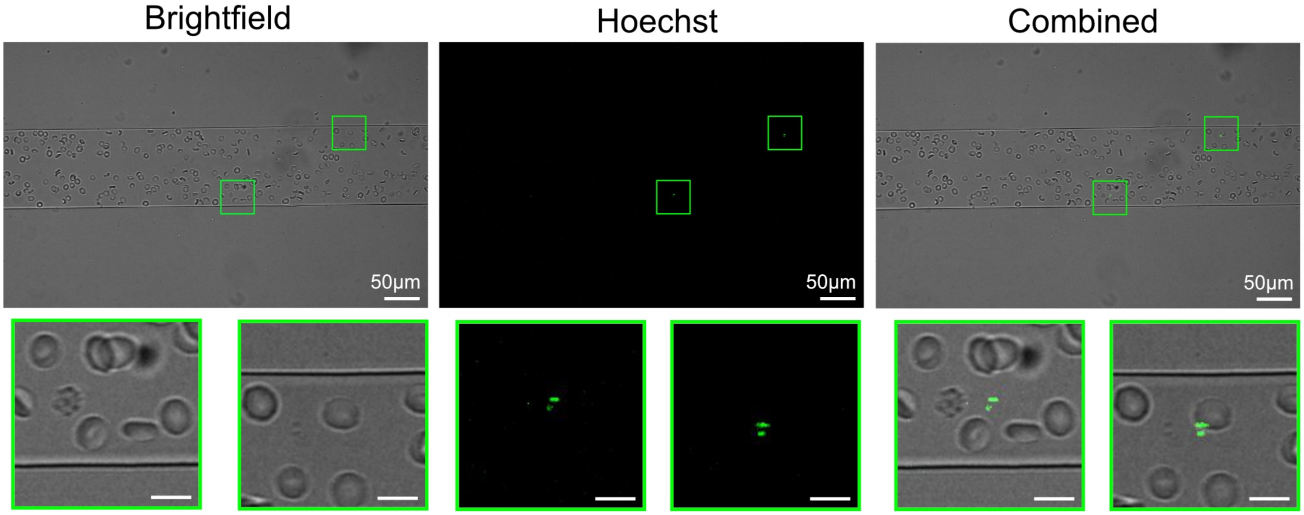


**Fig. S2. Comparison of infected erythrocytes in a microfluidics channel under flow with Hoechst 33342** Unless indicated otherwise, the scale bars are 10 $\mu$m. Images were taken with a 20x objective. Infected erythrocytes stained with Hoechst 33342 at 1 in 10000 concentration for 10 mins. The experiment was run with a single channel device with a 100 $\mu$m width and 10 $\mu$m depth. The green boxes contain at least one example of a ring (when the erythrocytes are followed across the channel, the fluorescent signal stays with them).

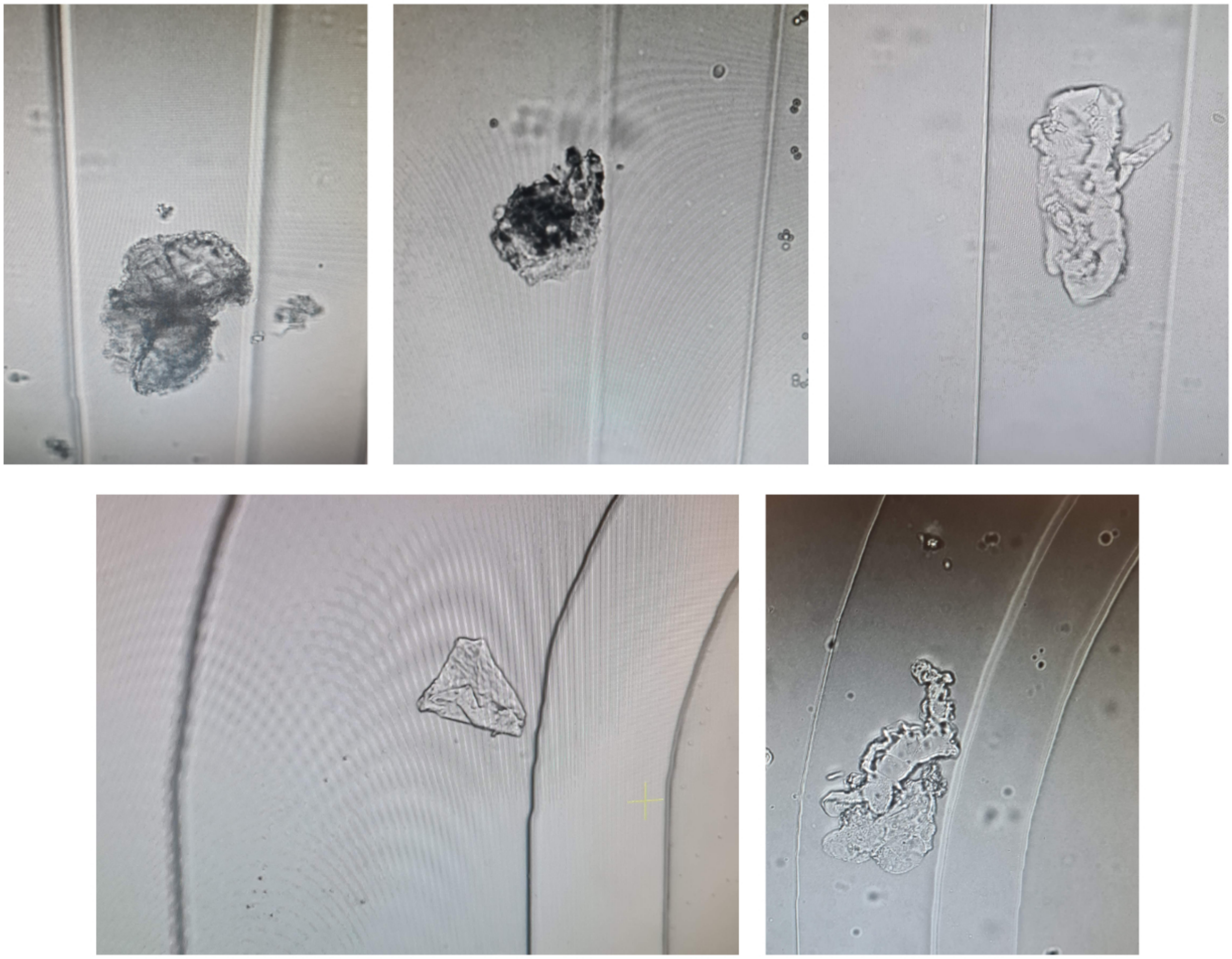

**Fig. S3.** Examples of non-cell debris seen in microfluidics channels.

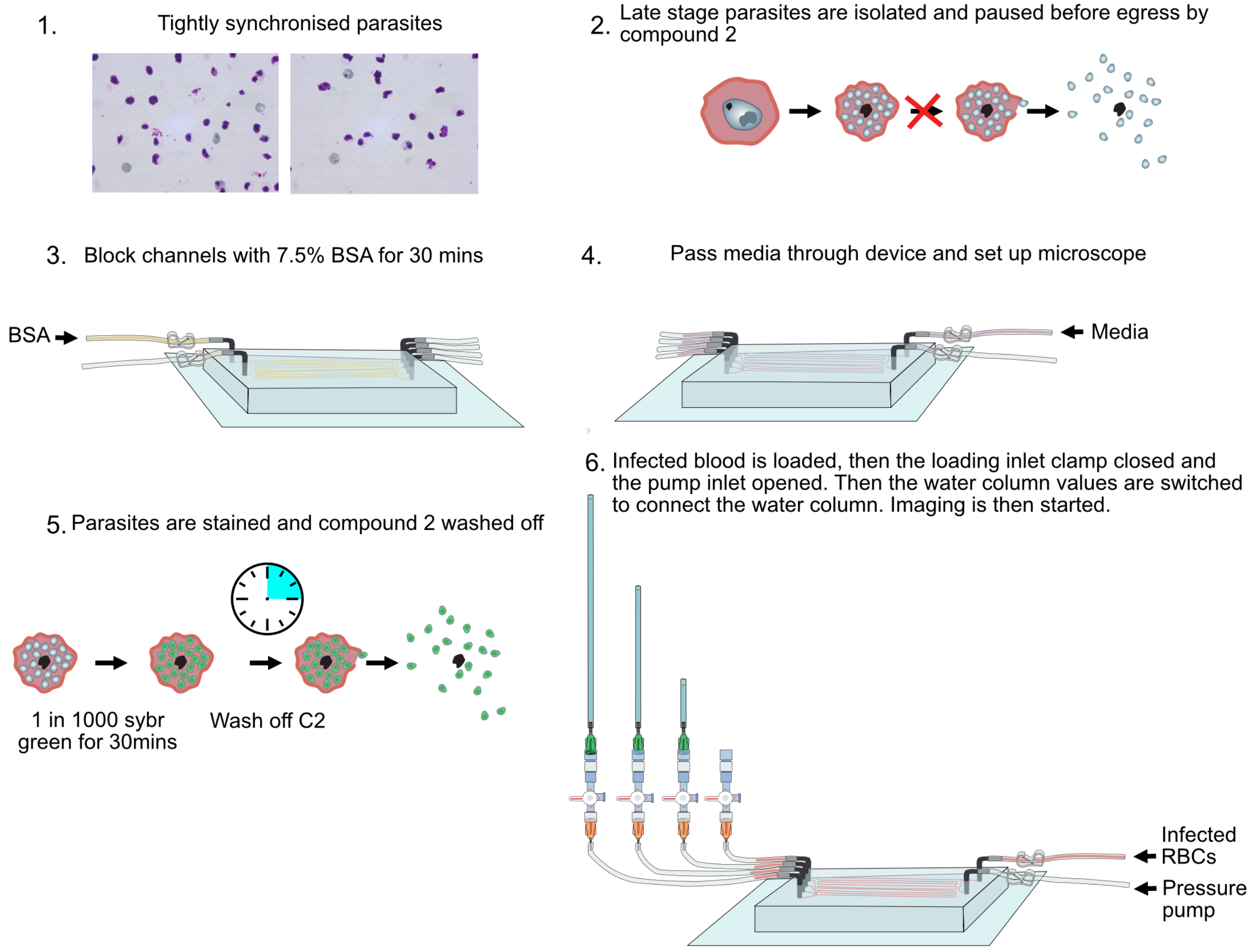


**Fig. S4. Summary of microfluidic experiment workflow** On the day of the experiment, Schizonts are isolated from a sample which has been tightly synchronised to the correct time window using Percol. These are then resuspended in compound 2 for 3-5.5 hours. The microfluidics chip is set up, and 7.5% BSA is passed into the channel and left for 30 mins at 37$^{o}$C. An aliquot of the sample is then pelleted and resuspended in sybr green and Compound 2 for 30 mins. The chip is then transferred to the microscope and connected to the inlet, outlet and pump tubes. The pump is turned on but the tube to it is clamped. Media is passed through the chip to confirm that there are no leaks; if any sign of detachment is detected, the chip is discarded. The chip is aligned, and the microscope focuses on the imaging field of view, ensuring that all channels are in focus. The Compound 2 is then washed off the parasite sample, blood is added to give the desired hematocrit and it is loaded into a syringe. This then gives a 10 min time window to load the sample into the chip via the inlet. The sample is loaded until it is seen in all the channels and ideally in the outlet tubing. The inlet is then clamped, the pump inlet tubing unclamped and then the valves to the water columns switched so the column is connected to the outlet.

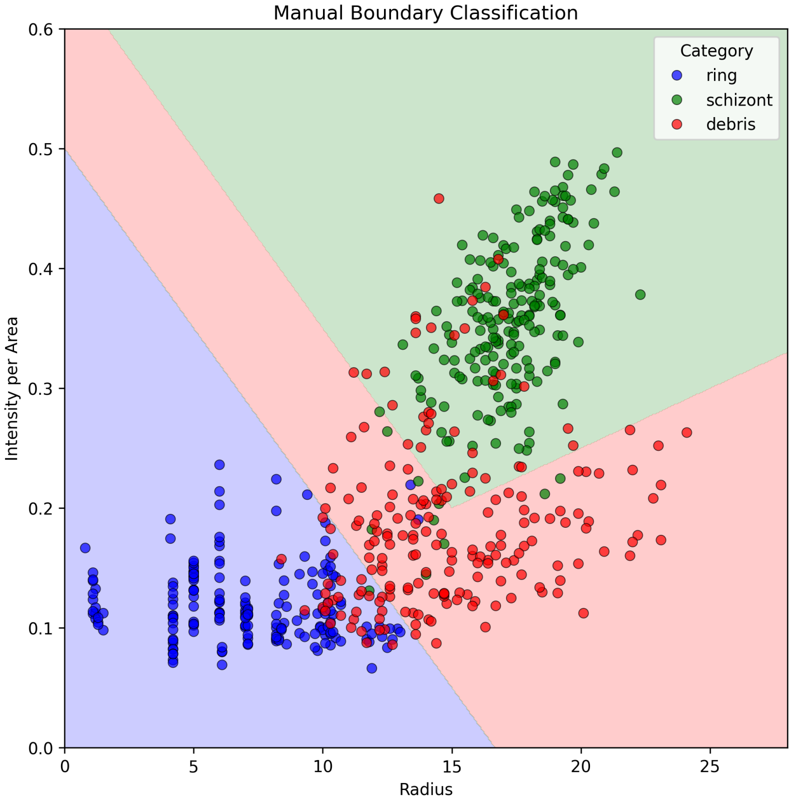


**Fig. S5.**

Graph to show how rings/merizoites, schizonts and debris were distinguished. Frames containing a cropped section of the brightfield and fluorescent image of different cells were exported from several videos. They were then manually classified as rings/merizoites, schizonts or debris. The intensity per area vs the radius of the fluorescent signal from the SYBR Green DNA dye for each cell was plotted. Boundaries were then defined between the three groups to maximise the correct classification of the three cell types. This is possible because rings/merizoites have a single copy of the genome, resulting in a small radius and a low-intensity signal. Schizonts have multiple copies of the genome tightly packed together, resulting in a fluorescent signal which is both larger in radius and more intense. Debris is likely made up of fragments of the infected RBC, stuck merizotites and/or hemozoin crystals. The radius and intensity of debris there tends to be between that of the rings/merizoites and schizonts.

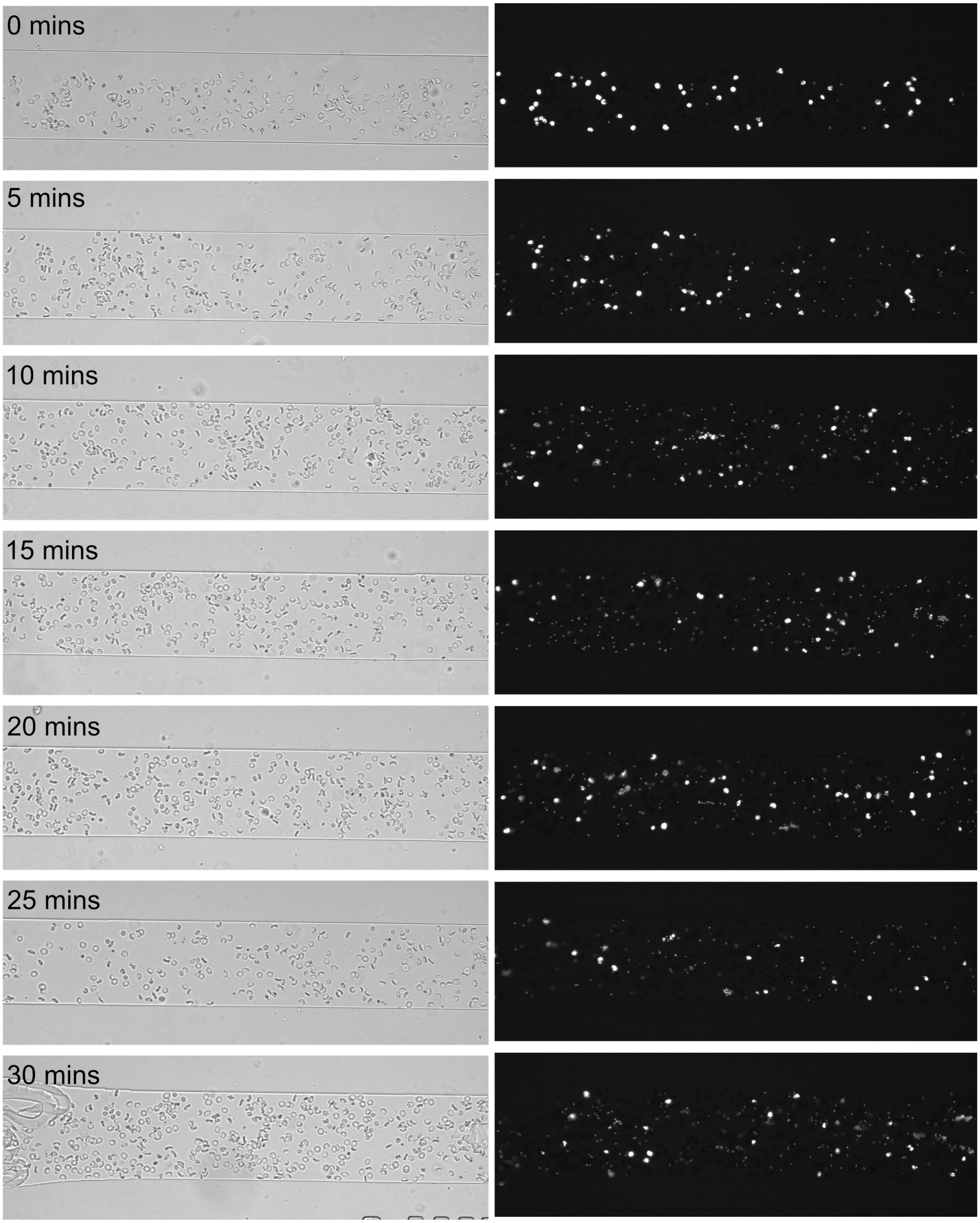


**Fig. S6. Example of frames taking over the course of an experiment to show the visual change in the proportion of different cell types.** Infected erythrocytes were stained with Sybr Green at a 1 in 5000 concentration for 45 mins. Time 0 is the time at which imaging began for this sample. Examples are shown of brightfield and fluorescent frames taken every 5 mins of the experimental run. The position of imaging was changed every 5 mins; the positions were determined based on the approximate flow rate of the cells so that roughly the same cells were imaged in every 5 mins segment. The experiment was run with a single channel device with a 100 $\mu$m width and 8.5 $\mu$m depth. The early images have a higher density of schizonts, then as time goes on, more merozoites are released as the schizonts egress. In the final image, there is a lower density of schizonts but a much higher density of small foci that indicate merozoite/rings.

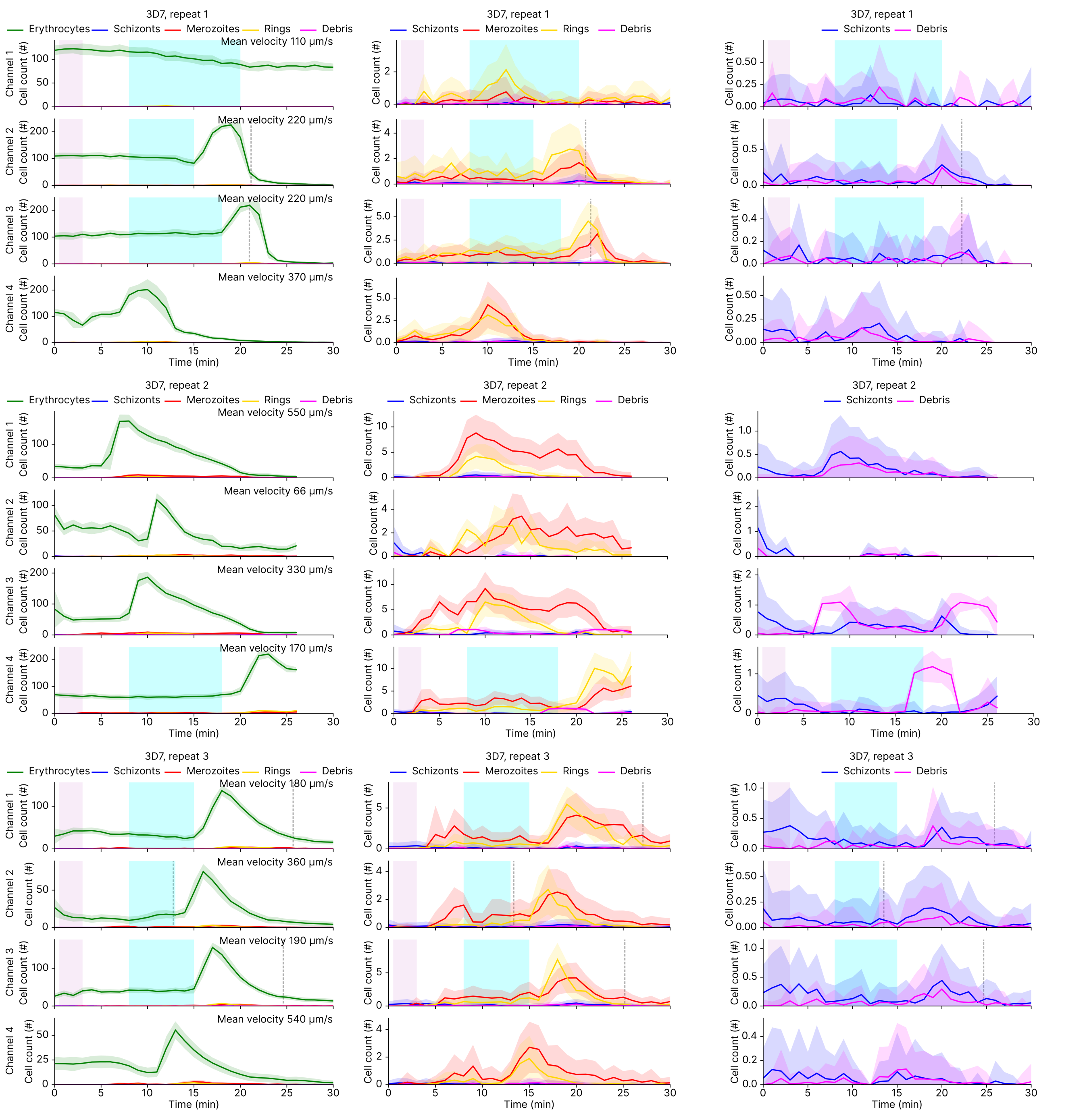
3D7, repeat 1
Erythrocytes
Schizonts
Merozoites
Rings
Debris
Mean velocity 110 μm/s
Mean velocity 220 μm/s
Mean velocity 220 μm/s
Mean velocity 370 μm/s
Channel 1
Channel 2
Channel 3
Channel 4
Cell count (#)
Time (min)
3D7, repeat 2
Mean velocity 550 μm/s
Mean velocity 66 μm/s
Mean velocity 330 μm/s
Mean velocity 170 μm/s
3D7, repeat 3
Mean velocity 180 μm/s
Mean velocity 360 μm/s
Mean velocity 190 μm/s
Mean velocity 540 μm/s

**S7.a**

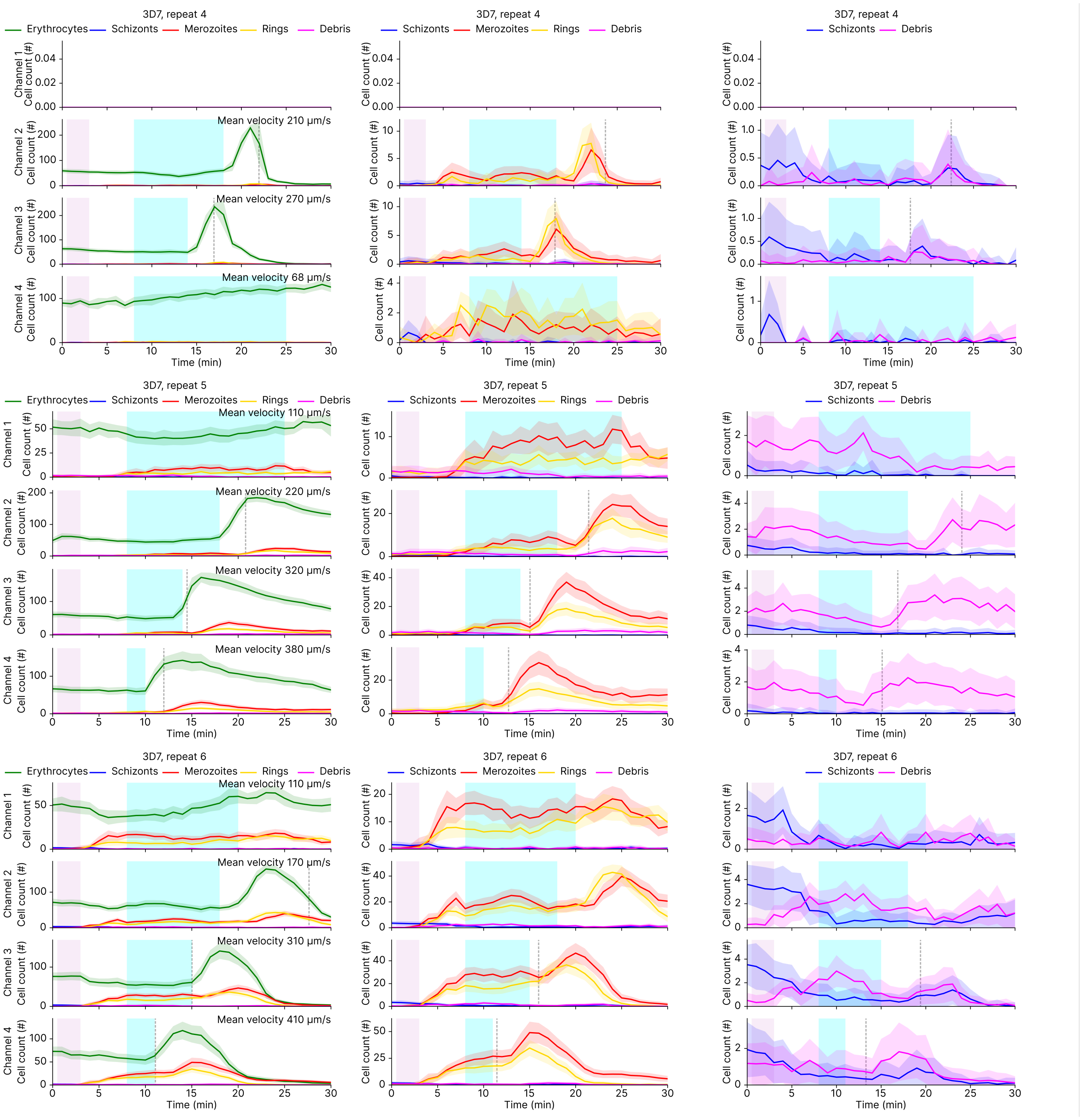
3D7, repeat 4
Erythrocytes
Schizonts
Merozoites
Rings
Debris
Channel 1
Channel 2
Channel 3
Channel 4
Cell count (#)
Mean velocity 210 µm/s
Mean velocity 270 µm/s
Mean velocity 68 µm/s
Time (min)
3D7, repeat 5
Mean velocity 110 µm/s
Mean velocity 220 µm/s
Mean velocity 320 µm/s
Mean velocity 380 µm/s
3D7, repeat 6
Mean velocity 110 µm/s
Mean velocity 170 µm/s
Mean velocity 310 µm/s
Mean velocity 410 µm/s

**S7.b**

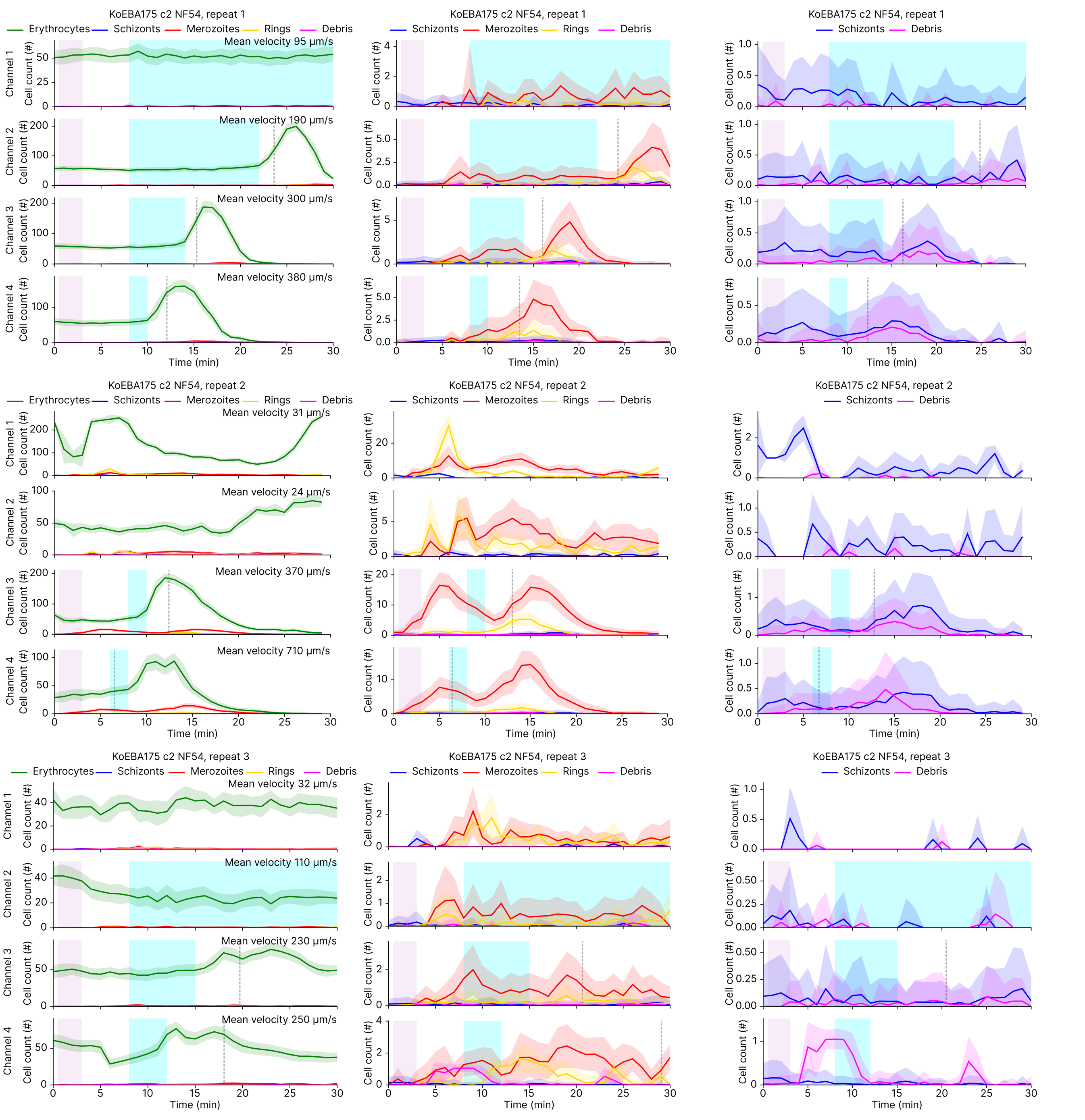
KoEBA175 c2 NF54, repeat 1
Erythrocytes
Schizonts
Merozoites
Rings
Debris
Mean velocity 95 µm/s
Mean velocity 190 µm/s
Mean velocity 300 µm/s
Mean velocity 380 µm/s
KoEBA175 c2 NF54, repeat 2
Mean velocity 31 µm/s
Mean velocity 24 µm/s
Mean velocity 370 µm/s
Mean velocity 710 µm/s
KoEBA175 c2 NF54, repeat 3
Mean velocity 32 µm/s
Mean velocity 110 µm/s
Mean velocity 230 µm/s
Mean velocity 250 µm/s
Channel 1
Channel 2
Channel 3
Channel 4
Cell count (#)
Time (min)

**S7.c**

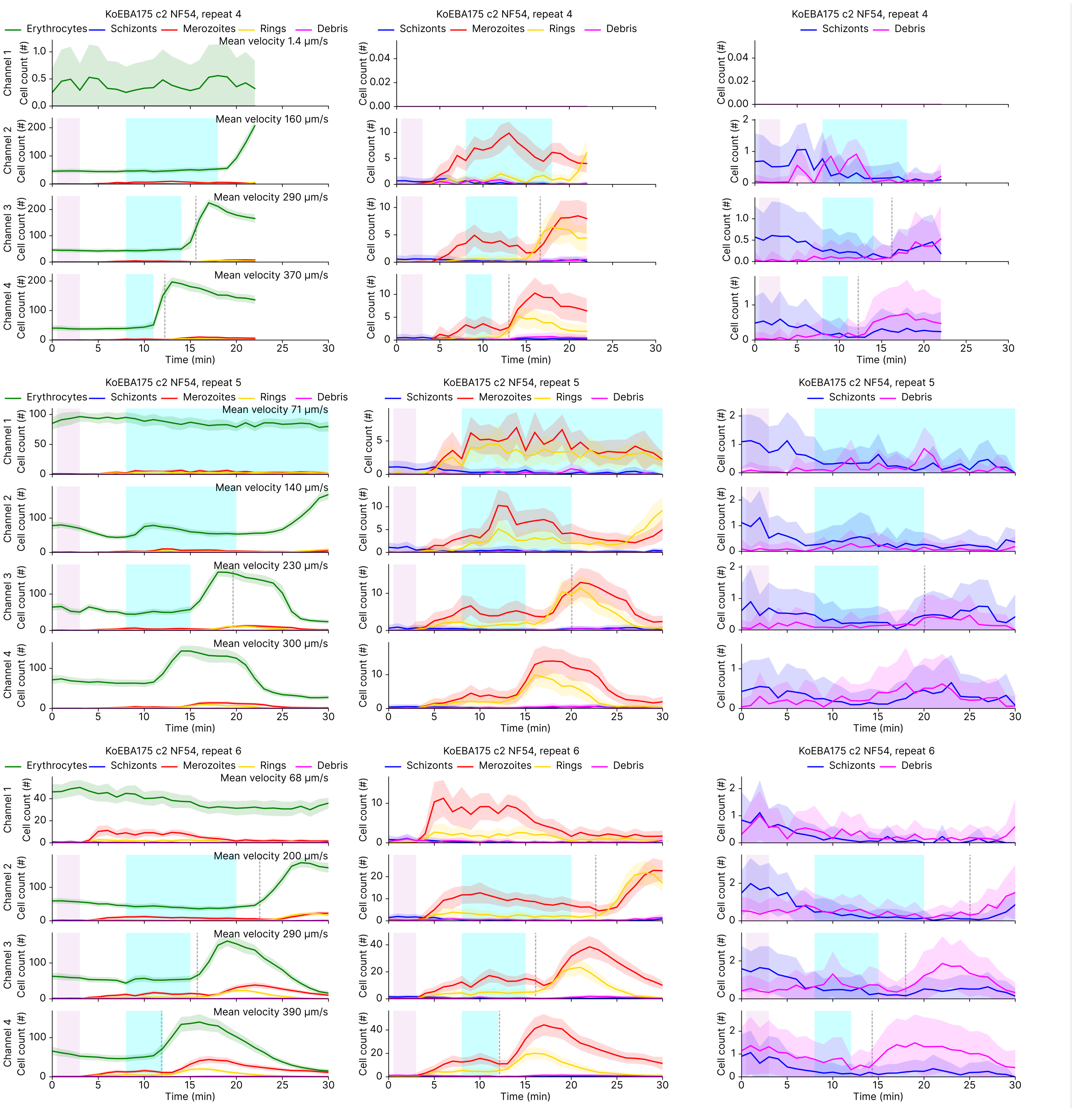
KoEBA175 c2 NF54, repeat 4
Erythrocytes
Schizonts
Merozoites
Rings
Debris
Mean velocity 1.4 μm/s
Mean velocity 160 μm/s
Mean velocity 290 μm/s
Mean velocity 370 μm/s
KoEBA175 c2 NF54, repeat 5
Mean velocity 71 μm/s
Mean velocity 140 μm/s
Mean velocity 230 μm/s
Mean velocity 300 μm/s
KoEBA175 c2 NF54, repeat 6
Mean velocity 68 μm/s
Mean velocity 200 μm/s
Mean velocity 290 μm/s
Mean velocity 390 μm/s
Channel 1
Channel 2
Channel 3
Channel 4
Cell count (#)
Time (min)

**S7.d**

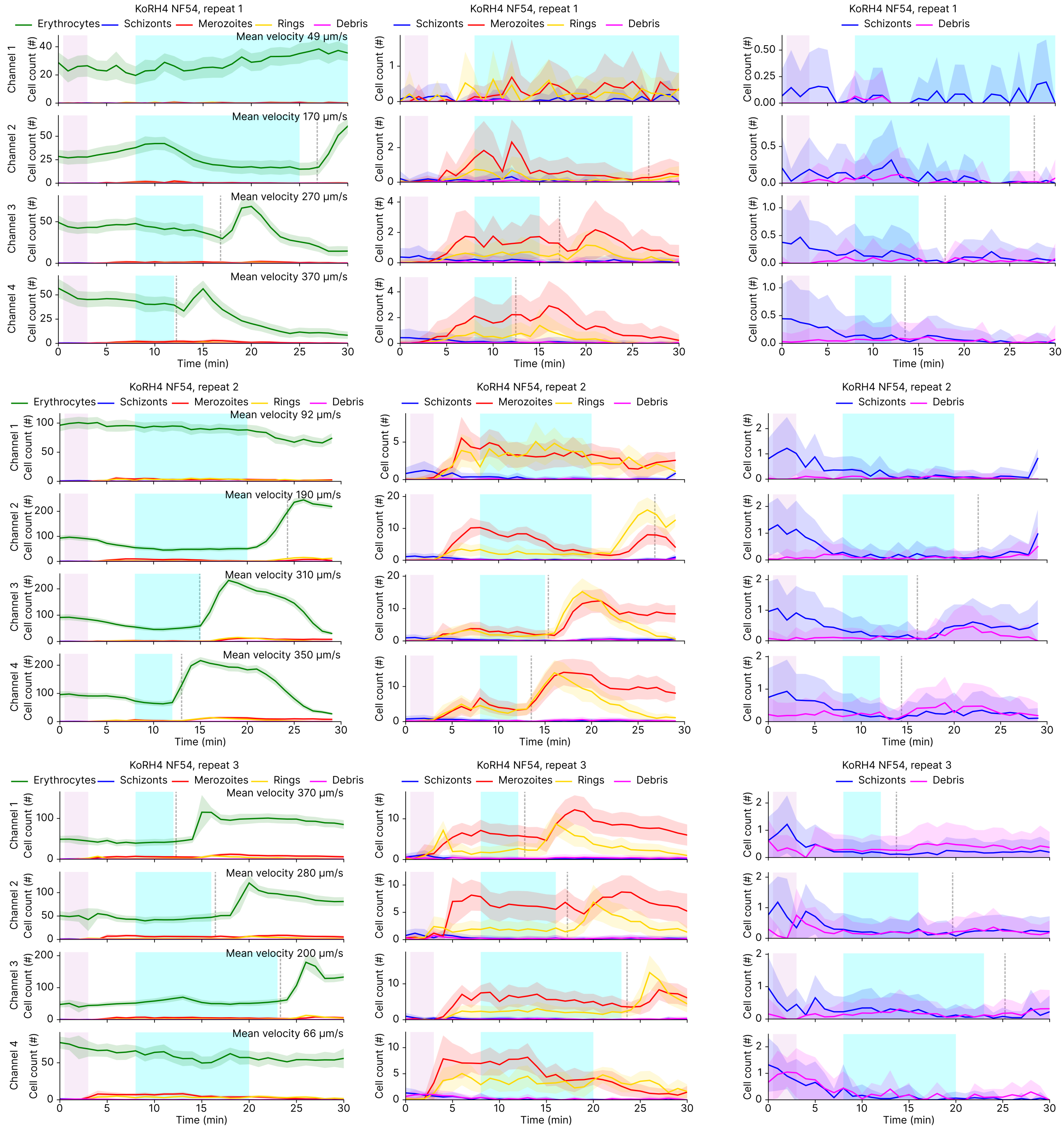
KoRH4 NF54, repeat 1
Erythrocytes
Schizonts
Merozoites
Rings
Debris
Mean velocity 49 µm/s
Mean velocity 170 µm/s
Mean velocity 270 µm/s
Mean velocity 370 µm/s
KoRH4 NF54, repeat 2
Mean velocity 92 µm/s
Mean velocity 190 µm/s
Mean velocity 310 µm/s
Mean velocity 350 µm/s
KoRH4 NF54, repeat 3
Mean velocity 370 µm/s
Mean velocity 280 µm/s
Mean velocity 200 µm/s
Mean velocity 66 µm/s
Channel 1
Channel 2
Channel 3
Channel 4
Cell count (#)
Time (min)

**S7.e**

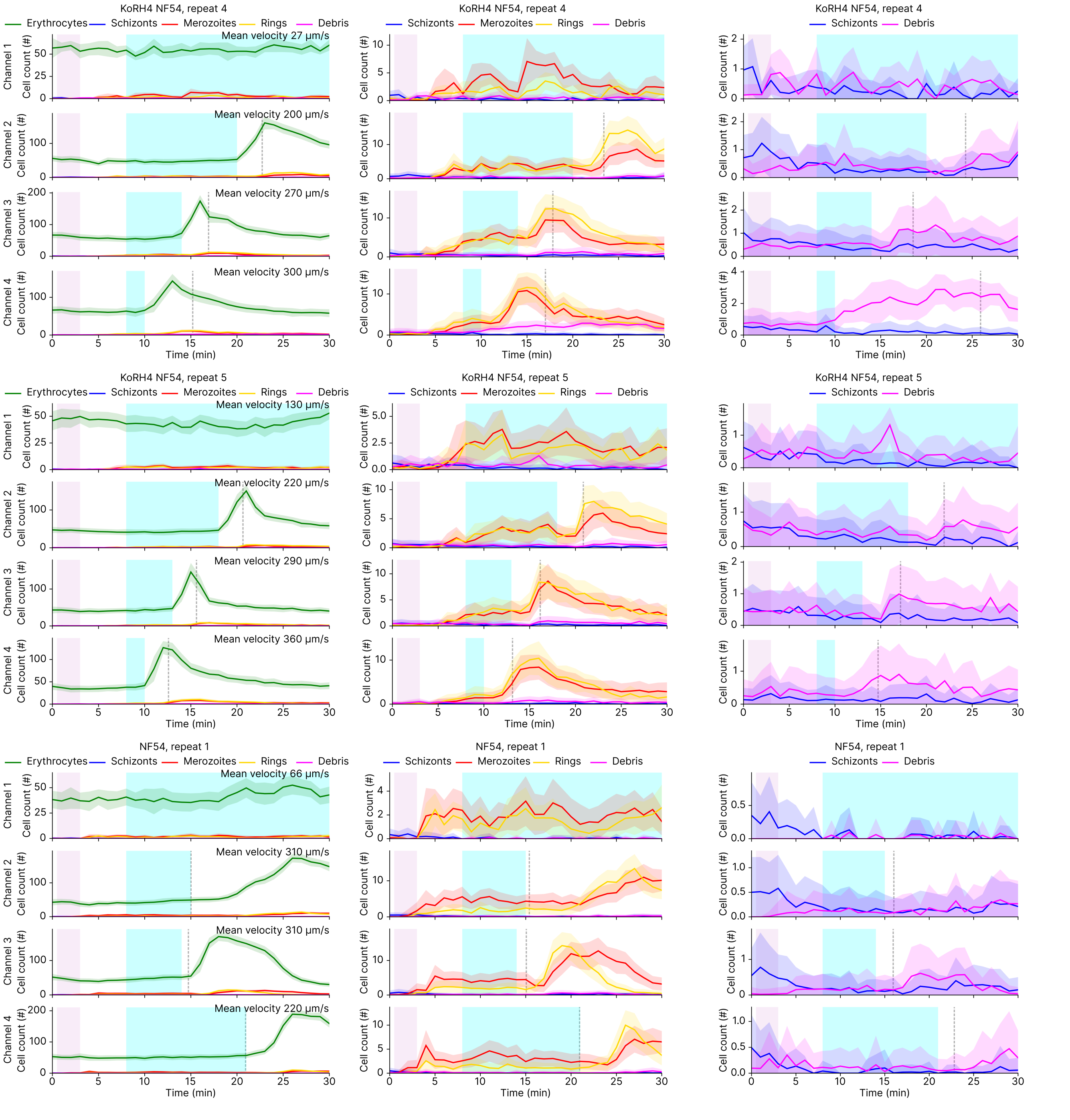
KoRH4 NF54, repeat 4
Erythrocytes Schizonts Merozoites Rings Debris
Schizonts Merozoites Rings Debris
Schizonts Debris
Mean velocity 27 μm/s
Mean velocity 200 μm/s
Mean velocity 270 μm/s
Mean velocity 300 μm/s
KoRH4 NF54, repeat 5
Mean velocity 130 μm/s
Mean velocity 220 μm/s
Mean velocity 290 μm/s
Mean velocity 360 μm/s
NF54, repeat 1
Mean velocity 66 μm/s
Mean velocity 310 μm/s
Mean velocity 310 μm/s
Mean velocity 220 μm/s
Channel 1
Channel 2
Channel 3
Channel 4
Cell count (#)
Time (min)

**S7.f**

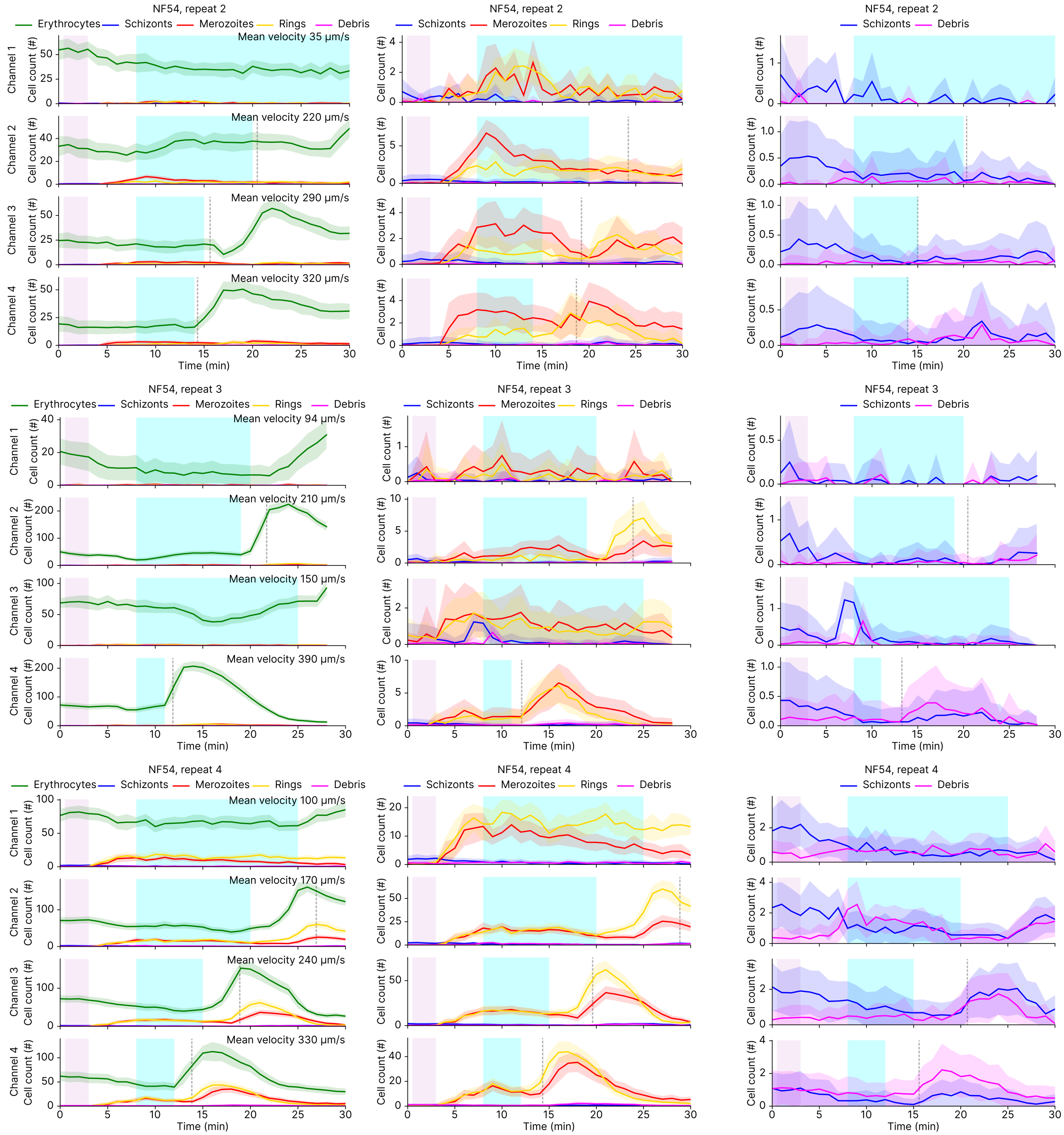
NF54, repeat 2
Erythrocytes
Schizonts
Merozoites
Rings
Debris
Mean velocity 35 µm/s
Mean velocity 220 µm/s
Mean velocity 290 µm/s
Mean velocity 320 µm/s
NF54, repeat 3
Mean velocity 94 µm/s
Mean velocity 210 µm/s
Mean velocity 150 µm/s
Mean velocity 390 µm/s
NF54, repeat 4
Mean velocity 100 µm/s
Mean velocity 170 µm/s
Mean velocity 240 µm/s
Mean velocity 330 µm/s
Channel 1
Channel 2
Channel 3
Channel 4
Cell count (#)
Time (min)

**S7.g**

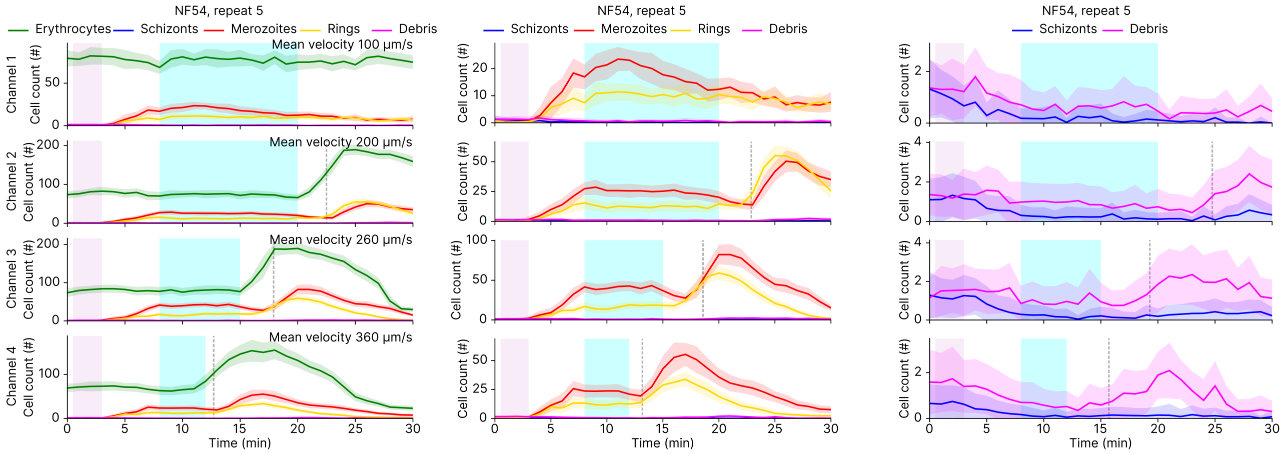


**S7.h**

**Fig. S7. Changes in cell counts of different cell types over time.** Plots show the change in different cell types' count over time for each experiment run. There are three plots for each experiment, each showing a different scale, so that the different cell types are clear.

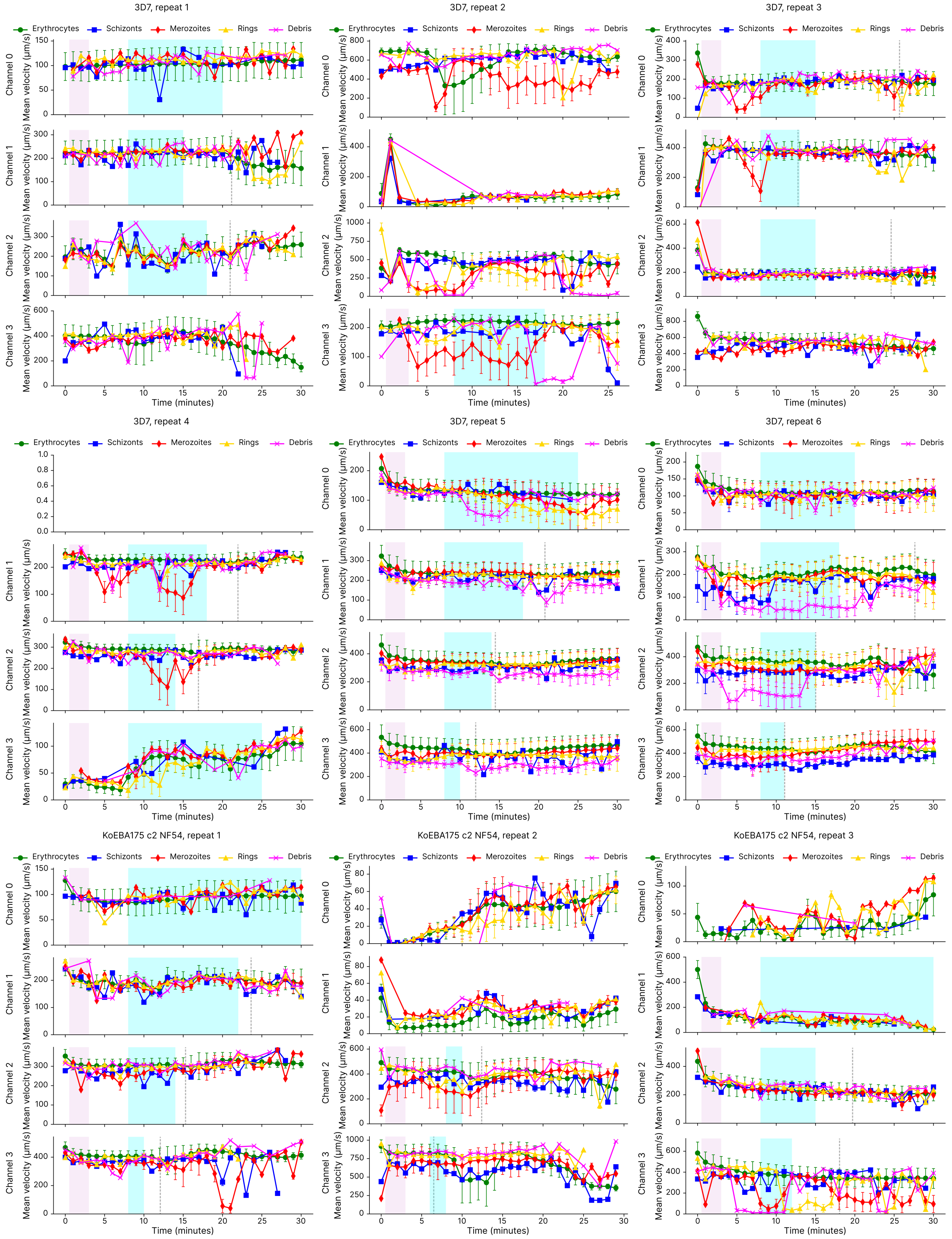
3D7, repeat 1
3D7, repeat 2
3D7, repeat 3
3D7, repeat 4
3D7, repeat 5
3D7, repeat 6
KoEBA175 c2 NF54, repeat 1
KoEBA175 c2 NF54, repeat 2
KoEBA175 c2 NF54, repeat 3
Erythrocytes
Schizonts
Merozoites
Rings
Debris
Channel 0
Channel 1
Channel 2
Channel 3
Mean velocity (μm/s)
Time (minutes)

S8.a

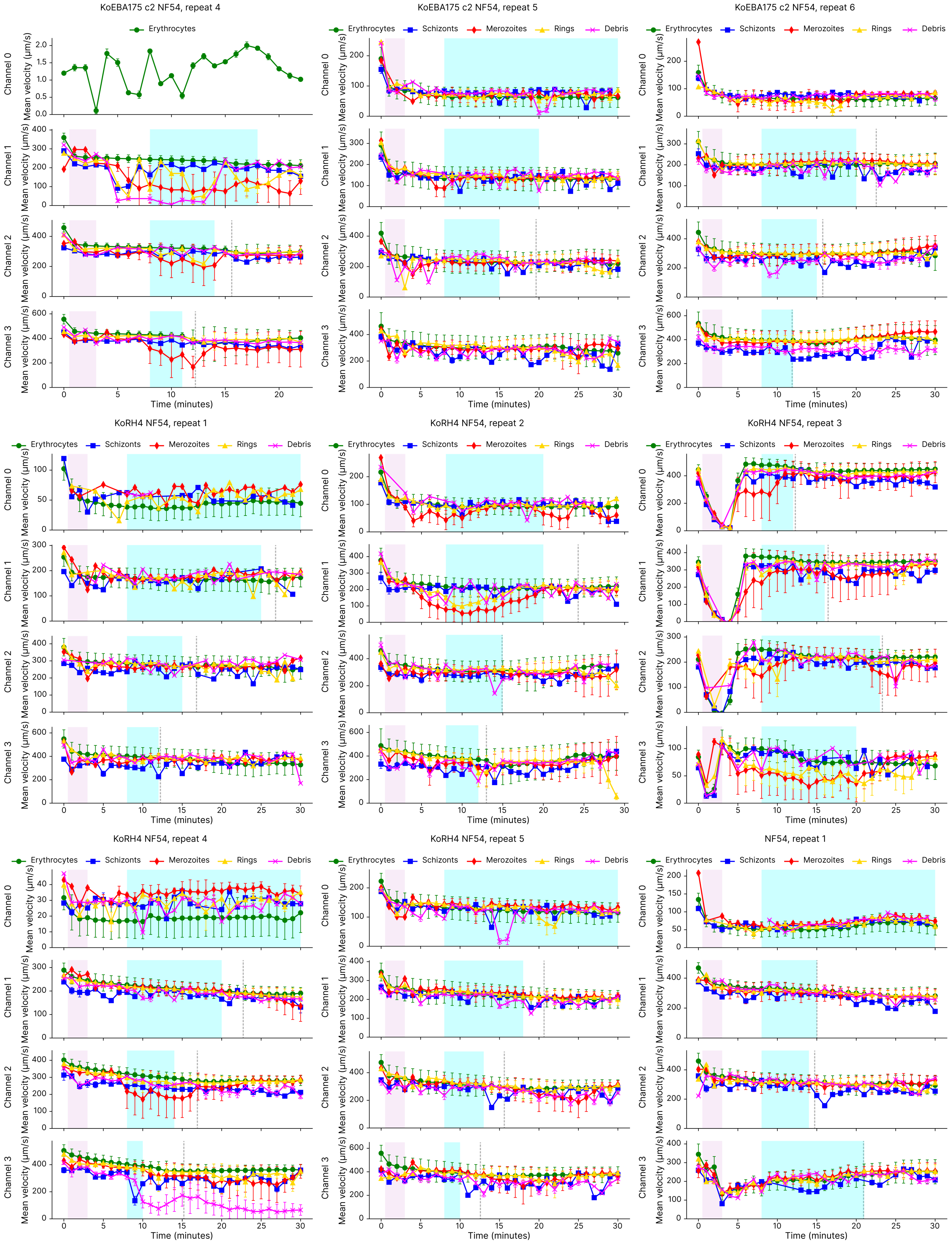
KoEBA175 c2 NF54, repeat 4
KoEBA175 c2 NF54, repeat 5
KoEBA175 c2 NF54, repeat 6
KoRH4 NF54, repeat 1
KoRH4 NF54, repeat 2
KoRH4 NF54, repeat 3
KoRH4 NF54, repeat 4
KoRH4 NF54, repeat 5
NF54, repeat 1
Erythrocytes
Schizonts
Merozoites
Rings
Debris
Channel 0
Channel 1
Channel 2
Channel 3
Mean velocity (μm/s)
Time (minutes)

S8.b

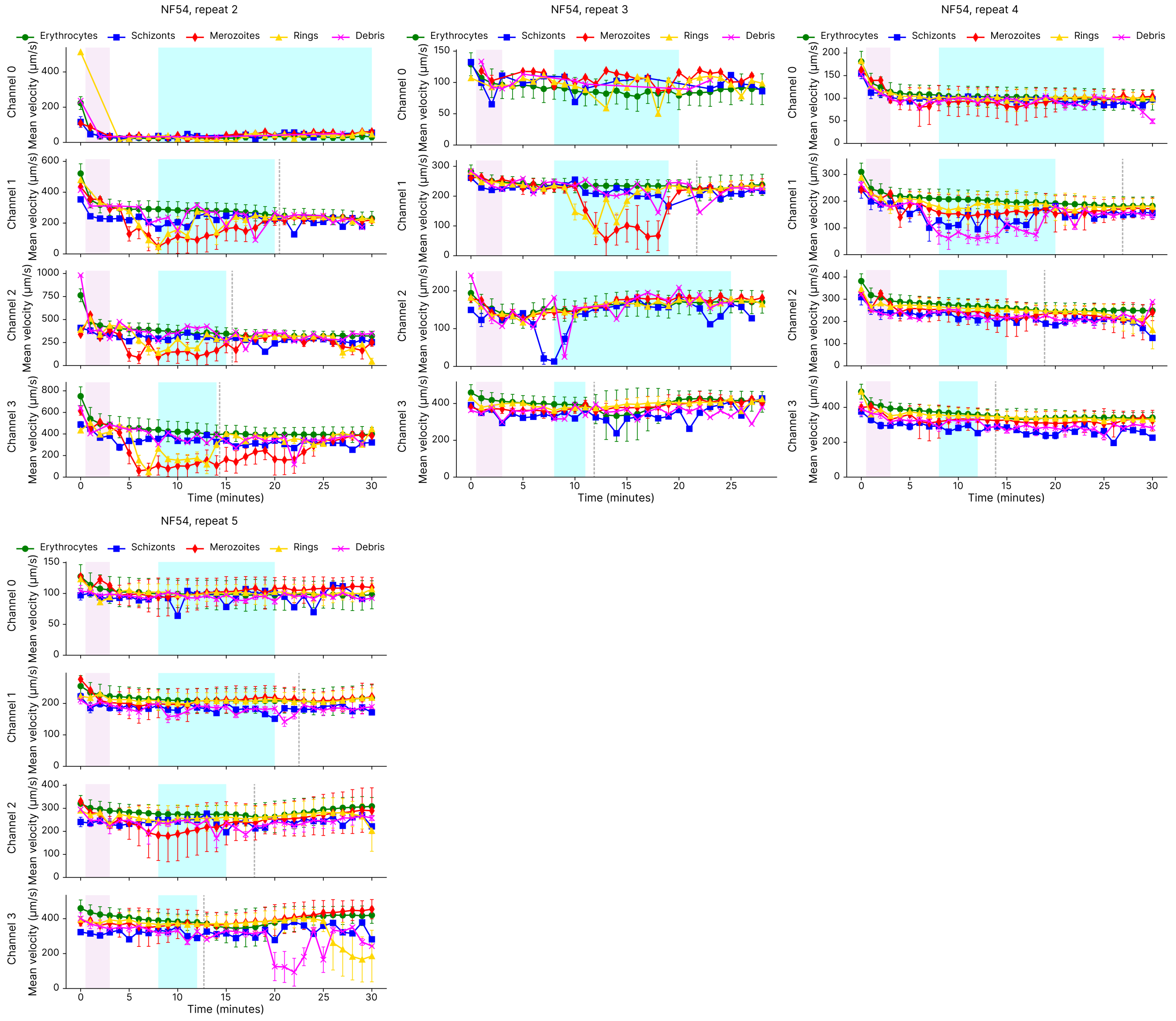


**S8.c**

**Fig. S8. Changes in flow rate of different cell types over time.**

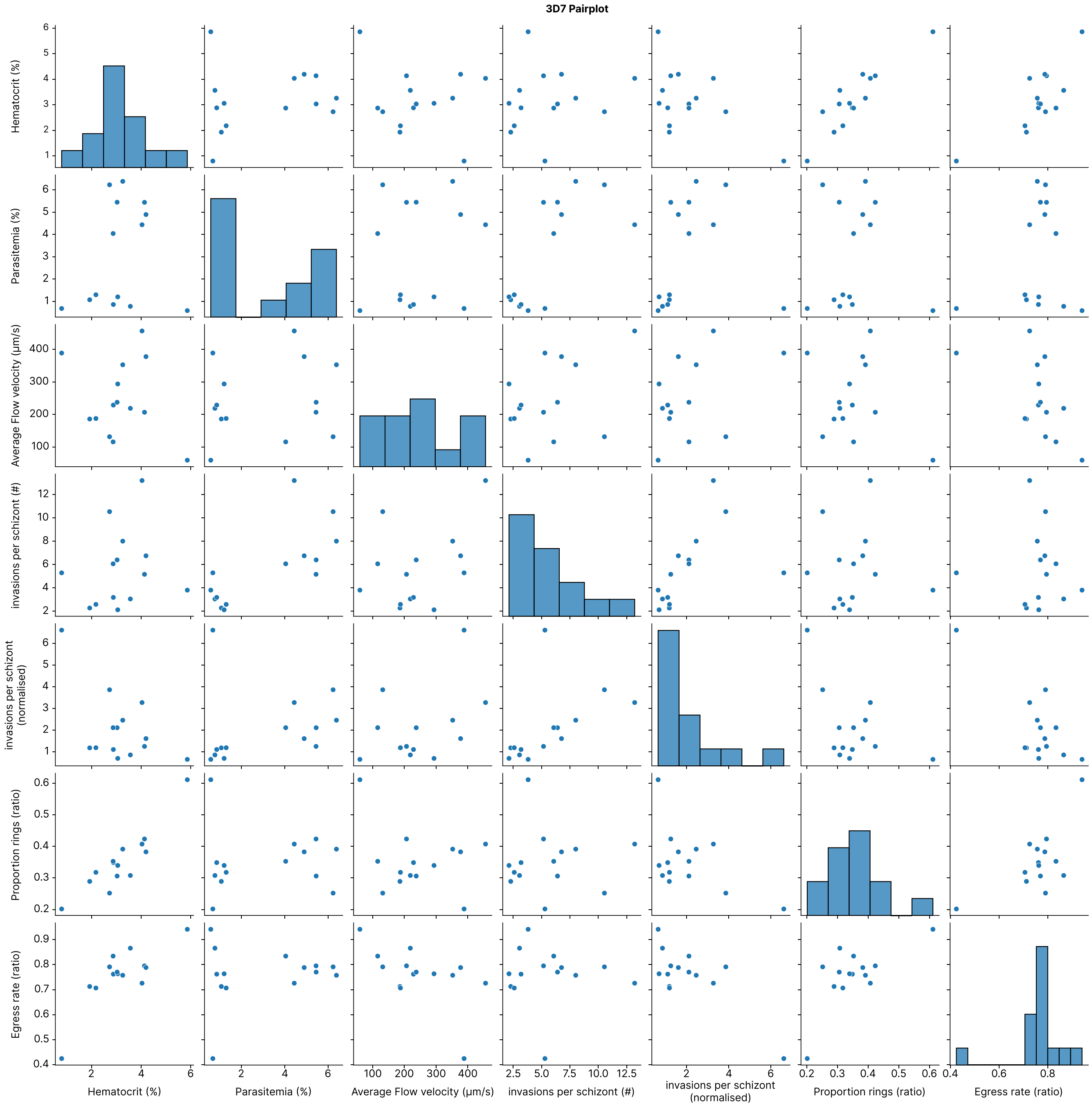
3D7 Pairplot
Hematocrit (%)
Parasitemia (%)
Average Flow velocity (µm/s)
invasions per schizont (#)
invasions per schizont (normalised)
Proportion rings (ratio)
Egress rate (ratio)

**S9.a**

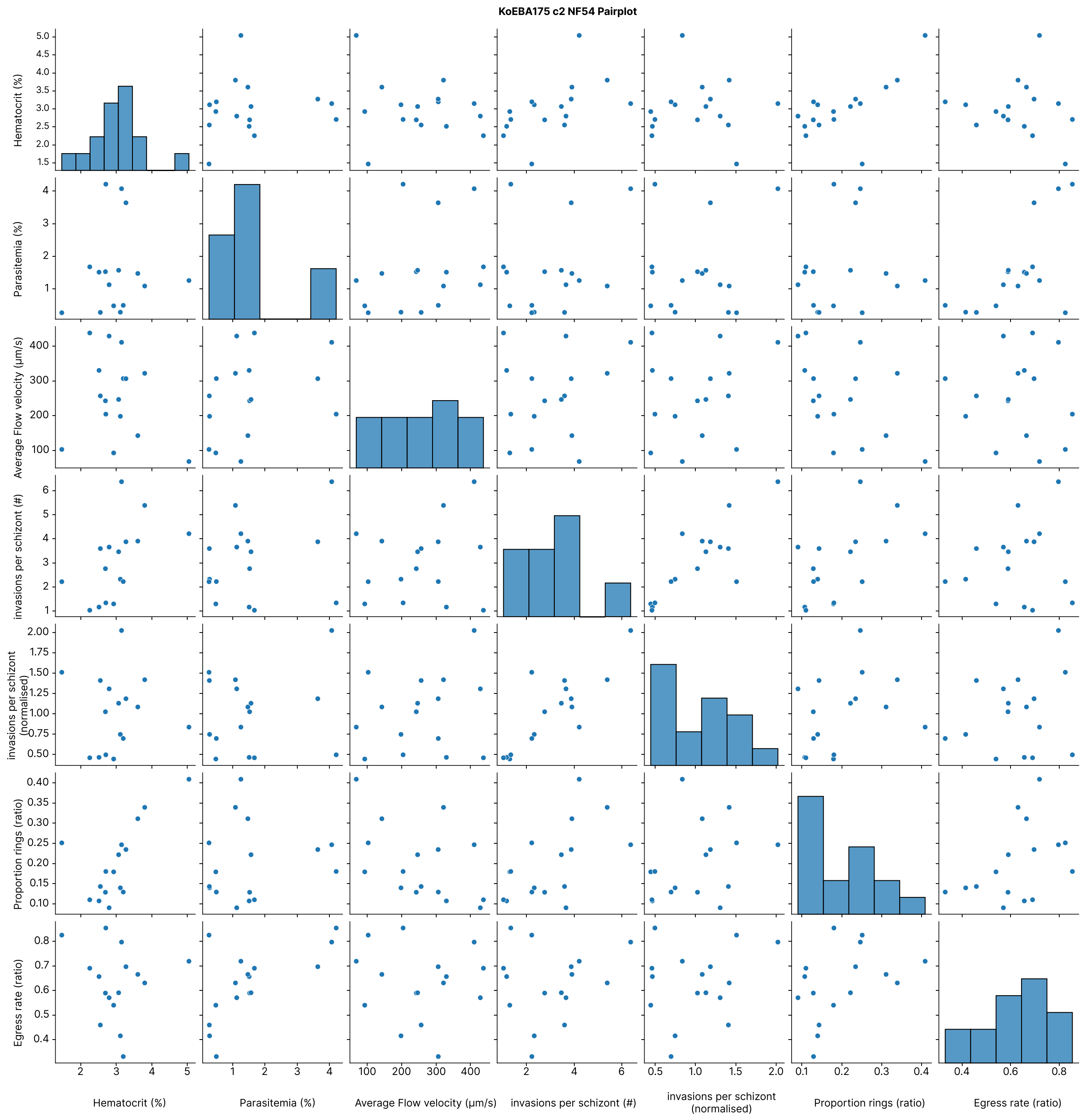
KoEBA175 c2 NF54 Pairplot
Hematocrit (%)
Parasitemia (%)
Average Flow velocity (μm/s)
invasions per schizont (#)
invasions per schizont (normalised)
Proportion rings (ratio)
Egress rate (ratio)

**S9.b**

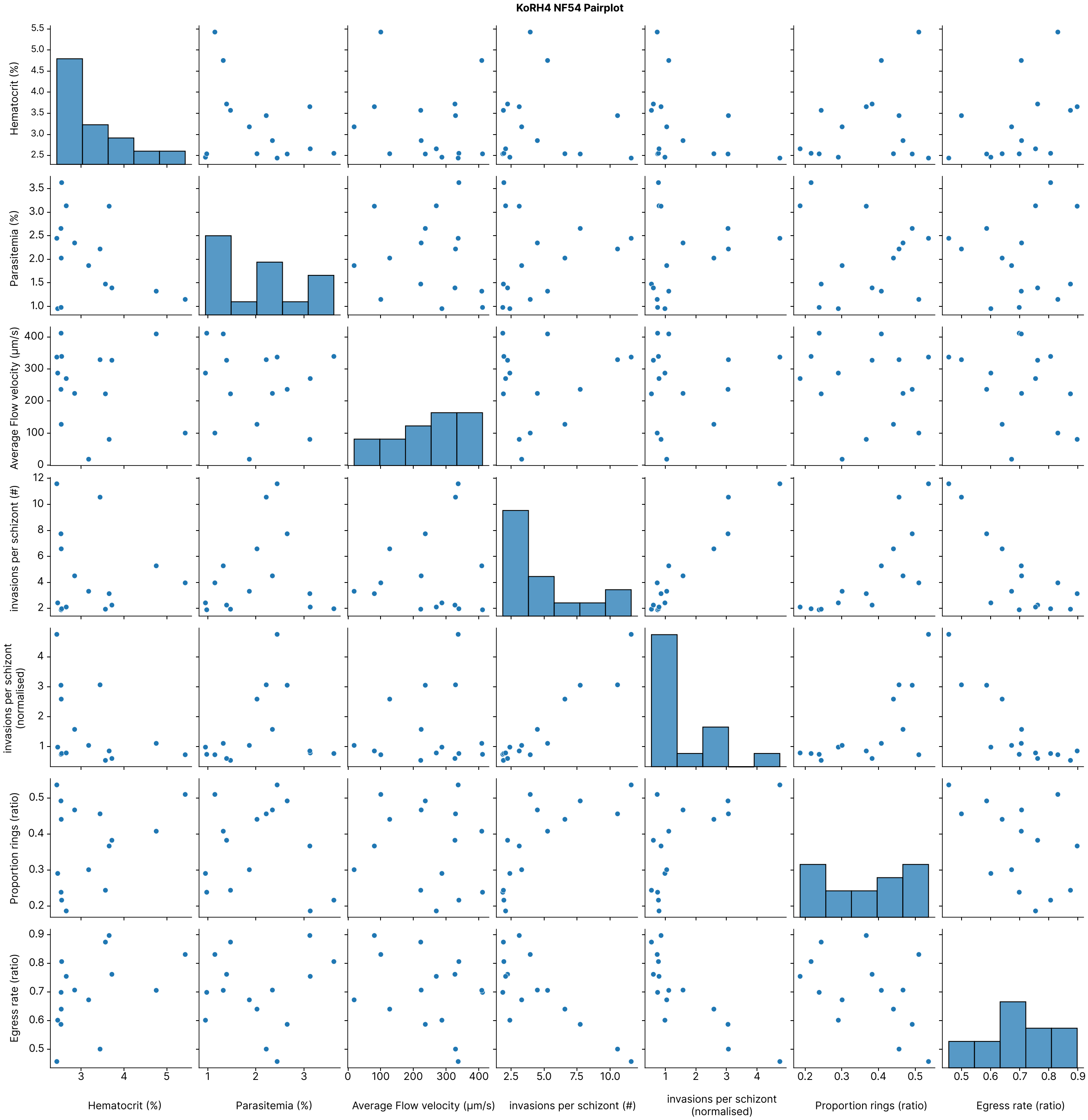

KoRH4 NF54 Pairplot
Hematocrit (%)
Parasitemia (%)
Average Flow velocity (µm/s)
invasions per schizont (#)
invasions per schizont (normalised)
Proportion rings (ratio)
Egress rate (ratio)


**S9.c**

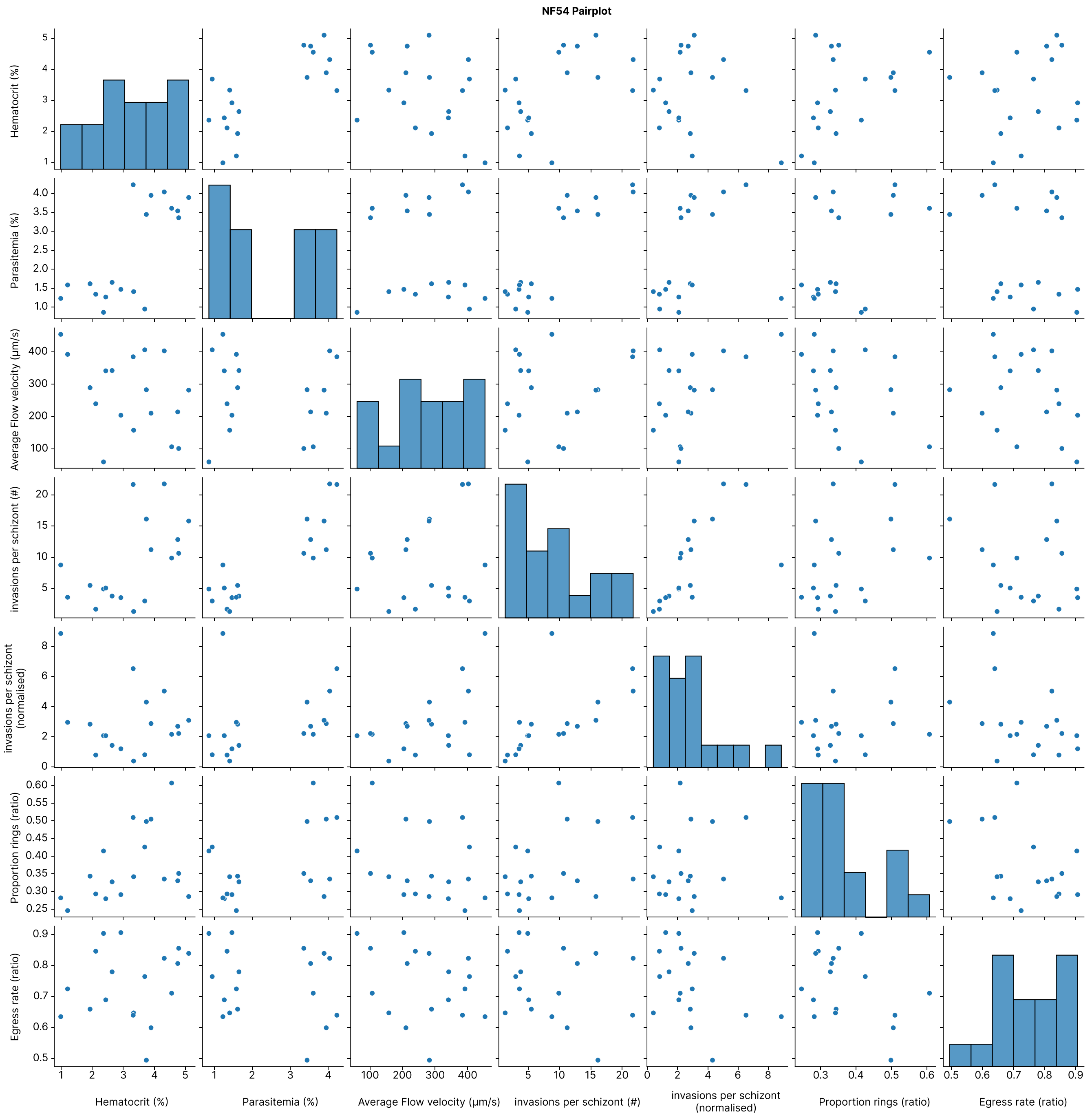


**S9.d**

**Fig. S9. Pairplots showing all combinations of variables considered for each species in turn.** Each point represents data from one channel and repeat.

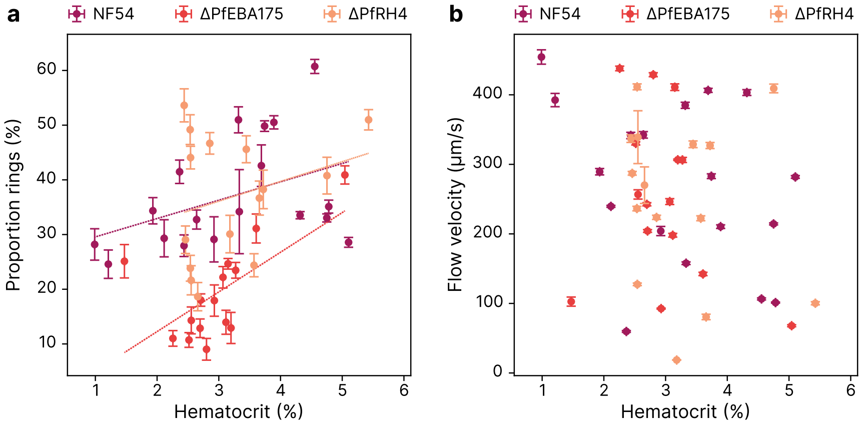


**Fig. S10.** Showing relationship between hematocrit, proportion rings and flow velocity for EBA175 and the two corresponding knock-out lines △PfEBA175 and △PfRH4. Each point represents data from one channel during one repeat of the experiment. Both A and B have error-bars in x and y representing standard deviation.
